# Spatial transcriptomics reveals a conserved segment polarity program that governs muscle patterning in *Nematostella vectensis*

**DOI:** 10.1101/2023.01.09.523347

**Authors:** Shuonan He, Wanqing Shao, Shiyuan (Cynthia) Chen, Ting Wang, Matthew C. Gibson

## Abstract

During early animal evolution, the emergence of axially-polarized segments was central to the diversification of complex bilaterian body plans. Nevertheless, precisely how and when segment polarity pathways arose remains obscure. Here we demonstrate the molecular basis for segment polarization in developing larvae of the pre-bilaterian sea anemone *Nematostella vectensis*. Utilizing spatial transcriptomics, we first constructed a 3-D gene expression atlas of developing larval segments. Capitalizing on accurate *in silico* predictions, we identified Lbx and Uncx, conserved homeodomain-containing genes that occupy opposing subsegmental domains under the control of both BMP signaling and the Hox-Gbx cascade. Functionally, *Lbx* mutagenesis eliminated all molecular evidence of segment polarization at larval stage and caused an aberrant mirror-symmetric pattern of retractor muscles in primary polyps. These results demonstrate the molecular basis for segment polarity in a pre-bilaterian animal, suggesting that polarized metameric structures were present in the Cnidaria-Bilateria common ancestor over 600 million years ago.

**Highlights:**

- *Nematostella* endomesodermal tissue forms metameric segments and displays a transcriptomic profile similar to that observed in bilaterian mesoderm
- Construction of a comprehensive 3-D gene expression atlas enables systematic dissection of segmental identity in endomesoderm
- *Lbx* and *Uncx*, two conserved homeobox-containing genes, establish segment polarity in *Nematostella*
- The Cnidarian-Bilaterian common ancestor likely possessed the genetic toolkit to generate polarized metameric structures

## Introduction

Bilaterian body plans are commonly constructed from a linear series of metameric segments along the anterior-posterior (AP) axis (Bateson, 1894; Christ et al., 1998; Diaz-Cuadros et al., 2021). A paraxial polarity program further subdivides each of these segments into opposing compartments with asymmetric developmental potential that will give rise to distinct parts of the nervous system, musculature and skeleton (Benazeraf and Pourquie, 2013; Onai et al., 2014). Segment polarization is thus a prominent feature observed in nearly all segmented animals and enables the formation of complex metameric structures with regionalized functions (Pourquie, 2000; Scott and Carroll, 1987). Still, despite superficial similarities, the genetic programs that establish segment polarity vary considerably, bringing into question the existence of an ancestral segment polarity program in the Urbilaterian common ancestor (Onai *et al*., 2014; Seaver, 2003; Tautz, 2004).

Cnidarians (jellyfish, hydroids, corals and sea anemones) are the sister group to bilaterians and are traditionally considered simple organisms with radial symmetry and an unsegmented body (Hyman, 1940). Contrary to this view, it has been recognized for over a century that many cnidarian species, especially in the basal class Anthozoa, display internal bilateral symmetry by forming metameric structures called gastric pouches along their directive axis (Pax, 1913). For instance, during larval development in the starlet sea anemone *Nematostella vectensis*, the endomesoderm undergoes tissue segregation to form eight bi-radially positioned segments (S1-S8, numbered clockwise with S1 being the largest), each constituting part of the future nerve net, musculature and gastrodermis (He et al., 2018; Leclere and Rentzsch, 2014; Steinmetz et al., 2017). During this process, pairs of nascent segment boundaries arise paraxially following a stereotypic sequence, regulated by a cnidarian Hox-Gbx code (He *et al*., 2018). On top of the metameric larval body plan, adult structures such as the retractor muscles are patterned in a segmentally-polarized manner, suggesting compartments with distinct developmental potential exist within each morphologically homogenous segment. Despite sharing similar design principles with bilaterian segments, little is known about the cellular and molecular basis for segment polarity establishment in cnidarians, hindering our ability to draw meaningful conclusions regarding the evolutionary origin of the metameric body plan.

The rapid development of spatial transcriptomics allows unbiased characterization of gene expression patterns in diverse biological contexts (Dries et al., 2021; Lohoff et al., 2022; Marx, 2021; van den Brink et al., 2020). However, widely-used spatial transcriptomic approaches such as Slide-seq are restricted to 2-dimensional tissue sections, offer limited resolution, and are difficult to employ in small 3-dimensional samples such as the embryos of marine invertebrates (Rodriques et al., 2019; Stickels et al., 2021). Fortunately, several computational approaches have been developed to circumvent these challenges, mostly relying on high-resolution landmark gene expression patterns to infer the likely coordinates of a given cell from conventional single cell RNA-seq (scRNA-seq) datasets (Achim et al., 2015; Deng et al., 2019; Karaiskos et al., 2017; Moriel et al., 2021; Nitzan et al., 2019; Satija et al., 2015).

Here we employed an *in silico* spatial transcriptomic approach to explore the molecular basis for segment polarity establishment in *Nematostella*. By generating a tissue-enriched scRNA-seq dataset, we first profiled the transcriptomic landscape of major endomesodermal cell clusters at mid-planula stage (72hpf), a time point at which the segmentation process has just completed. Next, guided by a set of 34 landmark genes, we constructed a 3-D spatial gene expression atlas (EndoAtlas) using the novoSpaRc algorithm, which predicts the spatial patterns for 15,542 genes that were expressed in the developing endomesoderm (Moriel *et al*., 2021; Nitzan *et al*., 2019). We validated our *in silico* predictions using fluorescent *in situ* hybridization (FISH) and systematically characterized segment identity markers downstream of the cnidarian Hox-Gbx hierarchy. Intriguingly, we also identified two conserved homeobox-containing genes, *Lbx* and *Uncx*, that establish opposing subsegmental domains concurrently with the segmentation process. Functional perturbations using short hairpin RNA (shRNA) knockdown and CRISPR/Cas9 mutagenesis demonstrated the requirements of the Bone Morphogenetic Protein (BMP) signaling pathway and the Hox-Gbx cascade in setting up the paraxial *Lbx-Uncx* polarity. In addition, *Lbx* loss of function abolished the molecular polarity and led to duplication of the retractor muscles in the resultant primary polyps. Lastly, phylogenetic analysis suggested that *Lbx* and *Uncx* first emerged in the Cnidaria-Bilateria common ancestor, consistent with an ancient origin for segment polarization in animals.

## Results

### Cellular and molecular profiling of developing endomesoderm in *Nematostella*

In *Nematostella*, the segmented larval body plan serves as a blueprint for the development of the adult anatomy. During metamorphosis, each segment takes on a different identity (tentacle-bearing or non-tentacle-bearing) while fusing with the neighboring segments at the boundaries to form the eight mesenteries which compartmentalize the gastric cavity of polyps (**Figure 1A**). Molecular markers for segmentally-polarized adult structures, such as the retractor muscles, start to exhibit asymmetric distribution at the planula stage, suggesting that distinct developmental identities are established early during the segmentation process (**Figure 1B**). These observations prompted us to focus on mid-planula stage larvae and investigate the molecular mechanisms for segment polarity.

**Figure 1.**
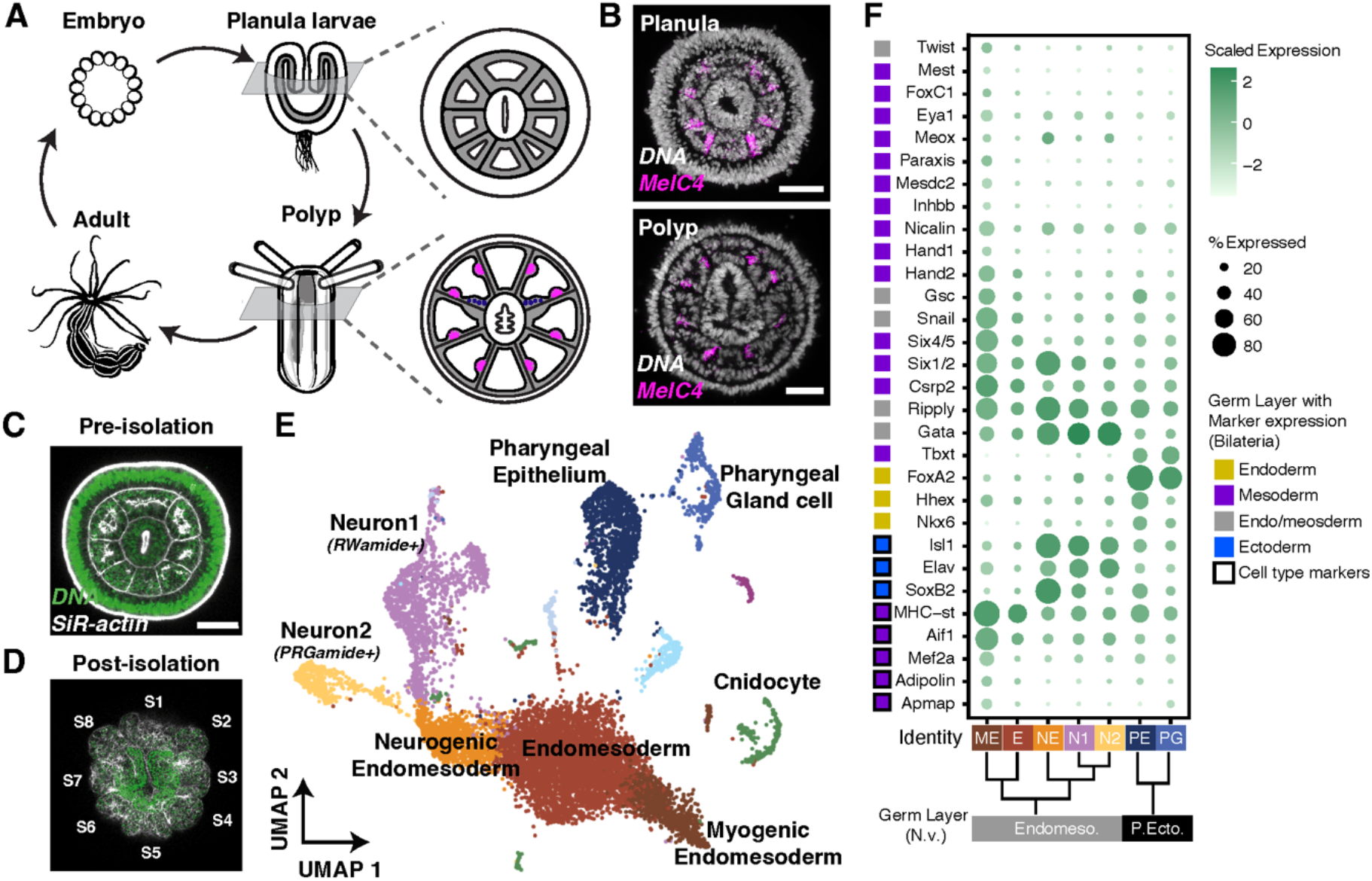
Molecular characterization of the developing *Nematostella* endomesoderm. (**A**) Schematic illustration of the *Nematostella* life cycle. At the polyp stage, certain endomesodermal cell types such as retractor muscle (*magenta*) and primordial germ cells (*dark blue*) are patterned in a segmentally polarized manner along the directive axis. (**B**) Fluorescent *in situ* hybridization showing the expression of *MHC-st*, a molecular marker for retractor muscles, at planula and polyp stages. The segmentally polarized expression of *MHC-st* is evident in the planula at 72hpf. Scale bars, 50μm. (**C** and **D**) F-actin and nuclear staining of 72hpf planula larvae before and after removal of the ectoderm. Tissue morphology was preserved during the process as all eight segments are clearly visible in endomesodermal isolates (S1 to S8). (**E**) UMAP projection of single cell RNA-seq data from endomesodermal isolates showing the different cell types identified. (**F**) Bubble plot illustrating the expression of homologs of bilaterian germ-layer and cell-type markers in different *Nematostella* cell types. Dendrogram in the bottom is based on the transcriptional similarities between different clusters. ME, myogenic endomesoderm; E, undifferentiated endomesoderm; NE, neurogenic endomesoderm; N1, neuron cluster 1; N2, neuron cluster 2; PE, pharyngeal ectoderm; PG, pharyngeal gland cells.

To analyze the cellular and molecular composition of larval segments on a global scale, we first employed scRNA-seq. At 72hpf, the endomesoderm constitutes a small proportion (<9%) of larval cells comparing to the ectoderm (Sebe-Pedros et al., 2018). We therefore combined osmolarity shock and chemical digestion to develop a tissue isolation method that specifically removes the ectoderm from developing larvae (**Figure 1C** and **D**, see also STAR Methods). Close examination of tissue isolates generated using this approach revealed the retention of part of the pharyngeal ectoderm adjacent to the intact segmented endomesoderm. By performing scRNA-seq on these isolates, we obtained 10,408 cells that passed a UMI cutoff of 500 (**Figure 1E,** see also STAR Methods). Subsequent UMAP projection identified 11 transcriptionally-distinct cell populations. Annotation of cluster identities was achieved using a set of published germ layer markers in *Nematostella*, such as *FoxA* (Pharyngeal ectoderm, PE) and *NvSnail1* (Endomesoderm, EN) (Fritzenwanker et al., 2004), as well as novel cluster markers identified by Seurat (**Figure S1**). Importantly, genes specifically expressed in the body wall ectoderm such as *Dlx* (Ryan et al., 2007) and *Anthox1* (Kamm et al., 2006; Ryan *et al*., 2007) were detected at minimal levels in this dataset, confirming the successful depletion of ectoderm during the isolation procedure (**Figure S1E**). As illustrated by the UMAP, EN is largely divided between the myogenic (ME) and neurogenic (NE) lineages, with the ladder further differentiating into two distinct larval neuron populations marked by the cnidarian neuropeptides *RWamide* (N1) and *PRGamide* (N2), respectively (**Figure S1G** and **H**) (Koch and Grimmelikhuijzen, 2019; Takahashi, 2020).

Interestingly, in *Nematostella* endomesoderm we observed enriched expression of genes whose homologs are typically associated with bilaterian mesoderm specification and differentiation, such as *FoxC1*, *Six1/2*, *Six4/5*, *Meox* and *Csrp2* (**Figure 1F**) (Chen et al., 2005; Mankoo et al., 2003; Miyasaka et al., 2007; Wilm et al., 2004). In addition, markers of bilaterian mesodermal cell types, including *MHC-st* (striated muscle), *Mef2a* (myocyte) and *Aif1* (macrophage/phagocyte) were enriched in the ME lineage (**Figure 1F, Figure S1I**) (Donovan et al., 2018; Nguyen et al., 1994; Noden et al., 1999). Conversely, pharyngeal ectodermal clusters were highly enriched for genes whose bilaterian homologs are involved in the induction of definitive endoderm, such as *FoxA2, Hhex* and *Nkx6* (**Figure 1F, Figure S1D**) (Burtscher and Lickert, 2009; Martinez Barbera et al., 2000; Pedersen et al., 2005). Taken together, these results demonstrate the complex cellular and molecular nature of the segmented *Nematostella* endomesoderm and argue for a revised view of germ-layer homology between cnidarians and bilaterians, consistent with previous candidate-centered gene expression studies (Martindale et al., 2004; Steinmetz, 2019; Steinmetz *et al*., 2017; Technau, 2020).

### *In silico* construction of a 3-dimensional gene expression atlas

To visualize endomesodermal gene expression on a global scale, we leveraged our single cell transcriptomic data to construct an *in silico* gene expression atlas. To achieve this, we applied novoSpaRc, a computational approach built on the premise that within a developing tissue cells with physical proximity tend to share similar transcription profiles, and thus one can infer the probable spatial distribution of individual cells within a defined space by solving an optimal-transport problem (Nitzan *et al*., 2019). Within the framework of novoSpaRc, we first generated a three-dimensional (3-D) model based on the morphological features of the actual endomesoderm (**Figure S2**) and projected cell clusters of endomesodermal origin (*Neuron1*, *Neuron2*, *Neuronal Precursor*, *Bulk Endomesoderm*, and *Myogenic Endomesoderm*, a total of 5,451 cells) onto 8,791 vertices located within this 3-D space (**Figure 2A**). In the initial unsupervised projection, novoSpaRc inferred the spatial position of cells based only on the internal structure of the data, and the prediction outcomes failed to recapitulate known gene expression patterns (exemplified by *Gdf5*, *Tbx15* and *Arp6*). This is likely because the default optimal arrangement of cells did not take into consideration the complex, metameric nature of the tissue, and indicated that spatial landmark genes were needed to guide the algorithm (Moriel *et al*., 2021). We therefore performed a fluorescent *in situ* hybridization (FISH) screen and identified 34 landmark genes that exhibited spatially distinct expression domains within the endomesoderm (**Figure S3A**). Binarized spatial expression matrices of these genes were then provided as additional input for novoSpaRc. The landmark-guided prediction successfully recapitulated the expression patterns of all 34 marker genes, with a mean Pearson correlation of 0.81 (**Figure S3B**).

**Figure 2.**
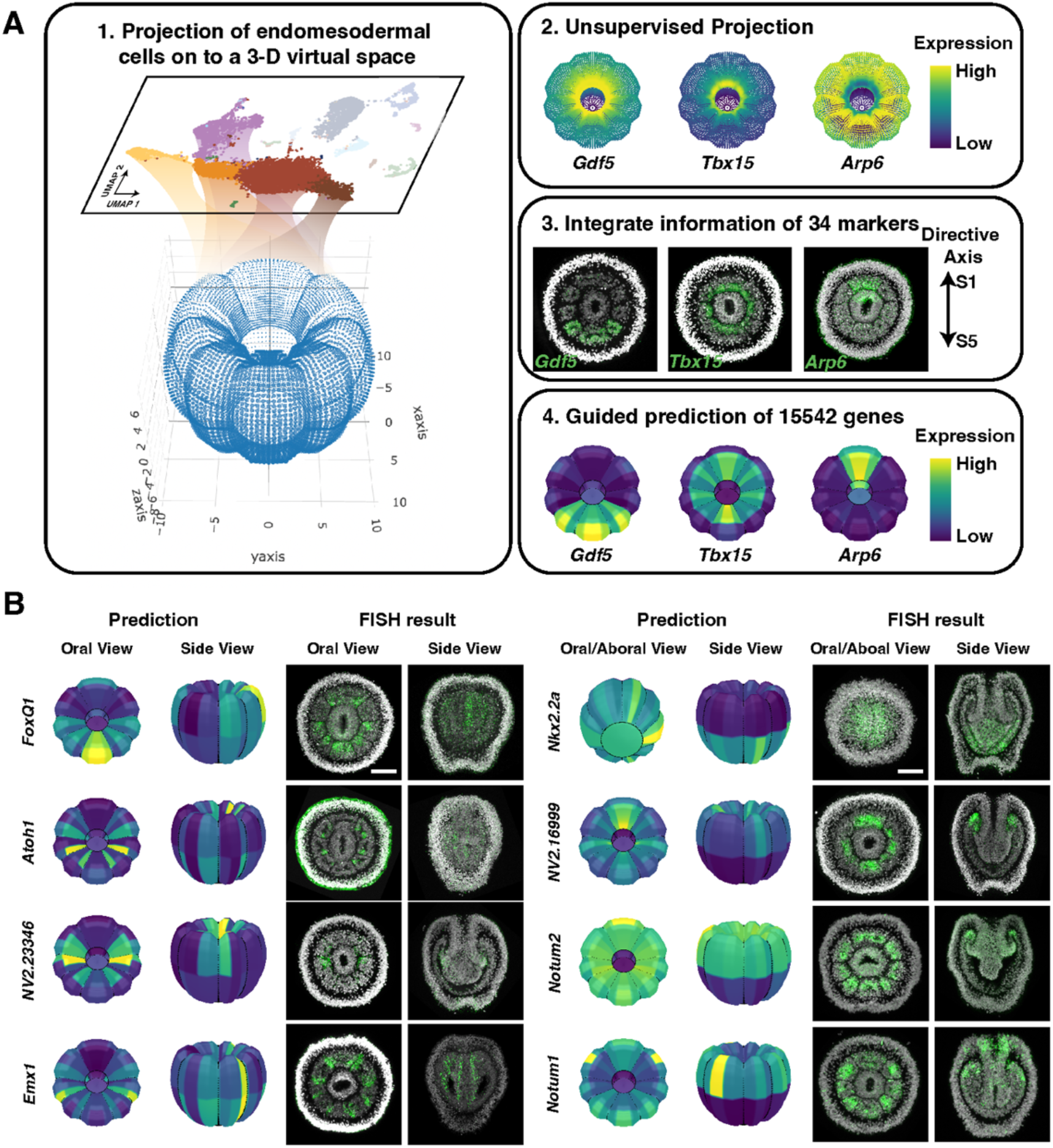
Construction of a 3D gene expression atlas in the developing *Nematostella* endomesoderm. (**A**) Workflow of Endo-atlas construction. From scRNA-seq data, cells of endomesodermal origin were projected on to a 3D virtual space that was based on the morphology of 72hpf planulae using novoSpaRc. A total of 34 landmark genes with distinct expression patterns were selected based on previous literature. Integrating binarized landmark gene expression patterns into novoSpaRc significantly increased the prediction accuracy. (**B**) Validation of in silico predictions of novel genes using FISH. Scale bar, 50μm.

We next validated the predictive power of Endo-atlas by analyzing the expression of 16 novel genes using FISH (**Figure 2B**). Among these we identified territorial genes expressed along the oral-aboral axis (e.g. *Notum1*, *Wntless*, *Nkx2.8*) as well as cell type-specific genes that displayed polarized expression in certain segments (*Atoh1* and *FoxQ1*). In all cases, the FISH results were in close accordance with our predictions, further supporting the accuracy of this approach. Lastly, we systematically predicted expression patterns for 15,542 genes that passed the minimum detection threshold in the *Nematostella* endomesoderm and constructed an online database named Endo-atlas (http://endoatlastest-env.eba-qtpfn7qz.us-east-1.elasticbeanstalk.com/).

### *Nematostella* endomesodermal segments possess distinct molecular identities

*Nematostella* endomesodermal segmentation is regulated by a group of Hox-Gbx genes expressed in an overlapping nested fashion (**Figure 3A**) (Chourrout et al., 2006; He *et al*., 2018; Hudry et al., 2014; Ryan *et al*., 2007). Guided by the Endo-atlas predictions, we next sought to identify segment identity genes acting downstream of the Hox-Gbx network to convey distinct developmental potential to otherwise identical segments. Indeed, we were able to confirm segment-restricted expression patterns of a large cohort of genes (**Figure 3B to E**). The polar segment S1 was marked by transmembrane receptors such as *Fgfr-like* and *CaSR*, with a subset of S1 cells adjacent to the pharyngeal ectoderm collectively expressing ADP-Ribosyltransferase *Art5*, neuropilin *Nrp2* and D1-dopamine receptor (**Figure 3B**). The enrichment of neuronal genes in segment S1 correlates with the asymmetric localization of *GLWamide*^+^ neurons within the developing endomesoderm (Watanabe et al., 2014). The tentacle-bearing segment pairs S2/S8 and S4/S6 shared several common markers, such as the structural protein *Col6A5*, the transcription factor *Zic3* as well as an unknown protein *Nv2.2940* (**Figure 3C**). Interestingly, only the S4/S6 segment pair expressed *Tgfr3, F5* and *Cdx* (**Figure 3D**), hinting at a molecular identity distinct from that of segments S2 and S6. Indeed, segments S2 and S8 give rise to exocoels in which incomplete mesenteries start to form during juvenile development, whereas S4 and S6 are endocoels that are incapable of secondary segmentation (Berking, 2007; Ikmi and Gibson, 2010; Ikmi et al., 2020). Segment S5, the polar segment opposite S1, possessed many unique molecular markers, such as the extracellular proteins *Efemp1* and *F8* as well as several Cnidaria-specific genes with unknown functions such as *Nv2.7863* (**Figure 3E**).

**Figure 3.**
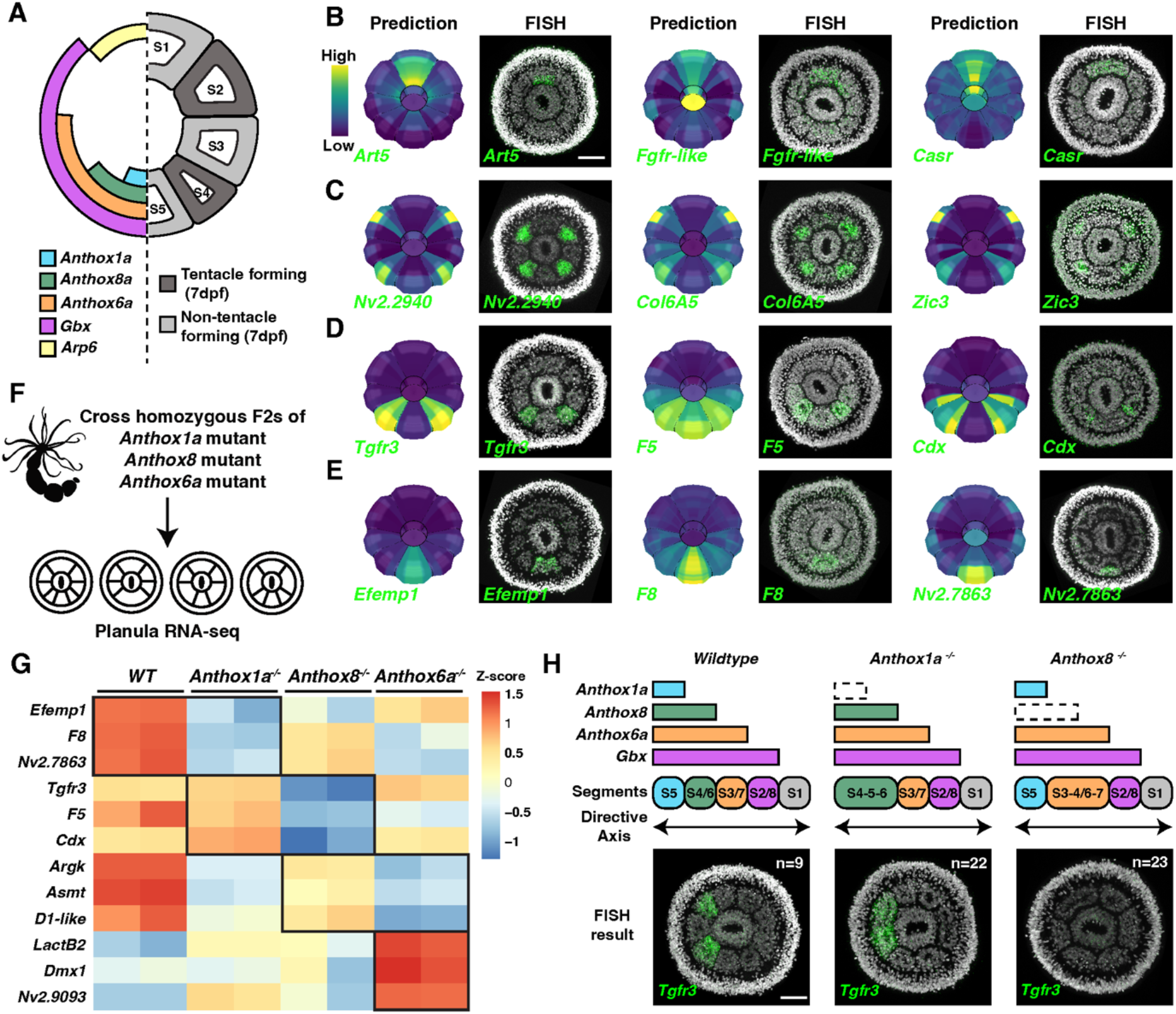
Characterization of segment identity genes in *Nematostella*. (**A**) Schematic of the endomesoderm demonstrating the Hox-Gbx molecular hierarchy underlying segment identity establishment. (**B-E**) Identified segment identity genes downstream of Hox-Gbx genes. (B) Segment S1 specific genes; (**C**) tentacle-bearing segment (S2/S4/S6/S8) specific genes; (**D**) S4/S6 specific genes; (**E**) Segment S5 specific genes. Color scale of *in silico* prediction indicates the relative expression level (*yellow*, high; blue, *low*). Scale bars, 50μm. (**F**) Experimental design of bulk RNA-seq under *Anthox1a*, *Anthox8a* and *Anthox6a* homozygous mutant backgrounds. (**G**) z-score of identified segmental identity genes in different mutant planulae (72hpf). Boxed up regions indicate expansion/retraction of segmental identities in different mutant backgrounds. (**H**) *Tgfr3* expression in different Hox mutants. Scale bars, 50μm.

To test whether Hox genes control the molecular identities of endomesodermal segments, we next performed RNA-seq in *Anthox1a, Anthox8* and *Anthox6a* homozygous mutant backgrounds. At 72hpf, the expression levels of the segment identity genes described above displayed drastic changes corresponding to the loss of each Hox gene (**Figure 3F**). For instance, in the absence of *Anthox1a* the physical boundaries of S5 are abolished, leading to the formation of a single fused segment S4-6 (He *et al*., 2018). Accompanying these morphological changes, we found that the expression of the S4/S6 identity marker *Tgfr3* expanded into the previous S5 territory, consistent with the expansion of S4/S6 identity across the fused segment (**Figure 3H**). Conversely, in the absence of *Anthox8* the S1-sided physical boundaries of S4/S6 are abolished, resulting in the formation of fused segments S3-4 and S6-7. In this case we observed the complete loss of *Tgfr3* expression (**Figure 3H**). These results are consistent with the altered tentacle patterns reported in Hox mutant polyps and demonstrate a molecular basis for homeotic transformation in *Nematostella*.

### *Lbx*, *Uncx* and the molecular polarity of *Nematostella* endomesodermal segments

Intriguingly, our Endo-atlas predictions identified two homeobox-containing transcription factors with segmentally polarized expression patterns: the *Ladybird* homolog *Lbx* and the *Unc-4* homolog *Uncx* (**Figure 4A** and **B**). In 72hpf planulae, *Lbx* expression specifically demarcated the S1-sided segment boundaries in segment pairs S2/S8, S3/S7, S4/S6 and was ubiquitously expressed in segment S5. *Uncx* was expressed in a complementary pattern to *Lbx*, where it was specifically enriched at the S5-sided boundaries in all three segment pairs and uniformly expressed in segment S1. *Lbx* and *Uncx* thus demarcate opposing territories with distinct molecular identities within all three segment pairs. Temporally, this polarity was established prior to the segmentation process, as *Lbx* stripes flanking future segment S1 and *Uncx* stripes flanking future segment S5 are detected before the formation of physical segment boundaries (**Figure S4**). These observations indicate that the *Lbx-Uncx* polarity program could be directly influenced by *Nematostella* Hox-Gbx genes, which also turn on prior to boundary establishment.

**Figure 4.**
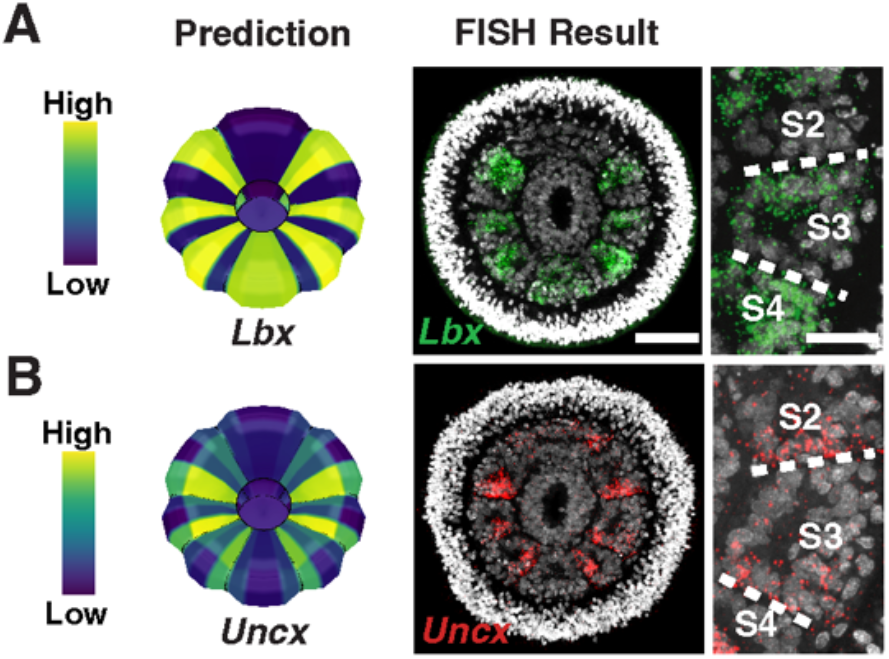
Identification of a segment polarity program downstream of Hox genes in *Nematostella*. (**A**) Endo-atlas prediction and FISH results demonstrating the polarized expression patterns of homeobox genes *Lbx* in 72hpf planula larvae. Scale bar, 50μm. Right panel shows the zoom in view of segment S3. Scale bar, 20 μm. (**B**) Endo-atlas prediction and FISH results demonstrating the polarized expression patterns of homeobox genes *Uncx* in 72hpf planula larvae. Right panel shows the zoom in view of segment S3.

To interrogate the upstream control of segment polarization, we examined the expression of *Lbx* and *Uncx* in *Anthox1a*, *Anthox8* and *Anthox6a* mutants (**Figure 5A** to **D**). *Anthox1a* mutants fail to form the S5 segment boundaries and instead generate an enlarged fusion segment S4-5-6. Consequently, the *Lbx* expression domain corresponding to segment S5 was absent, resulting in ubiquitous *Uncx* expressed in the center of the fusion segment (**Figure 5B**). *Anthox8* mutants fail to form boundaries between segments S3/S4 and S6/S7 and instead generate two enlarged fusion segments S3-4 and S6-7. Consequently, S1-sided *Lbx* stripes within segments S4 and S6 were abolished (**Figure 5C**). Both fusion segments still retained an *Lbx-Uncx* polarity along the directive axis, despite the increased segment size. Similarly, *Anthox6a* mutants fail to form boundaries between segments S2/S3 and S7/S8 and instead generate two enlarged fusion segments S2-3 and S7-8. Consequently, S1-sided *Lbx* stripes within segments S3 and S7 were abolished, and a positionally-shifted *Lbx-Uncx* polarity can be observed in both fusion segments (**Figure 5D**). Taken together, these results suggest that Hox genes provide segment-specific regulatory inputs towards *Lbx-Uncx* polarization.

**Figure 5.**
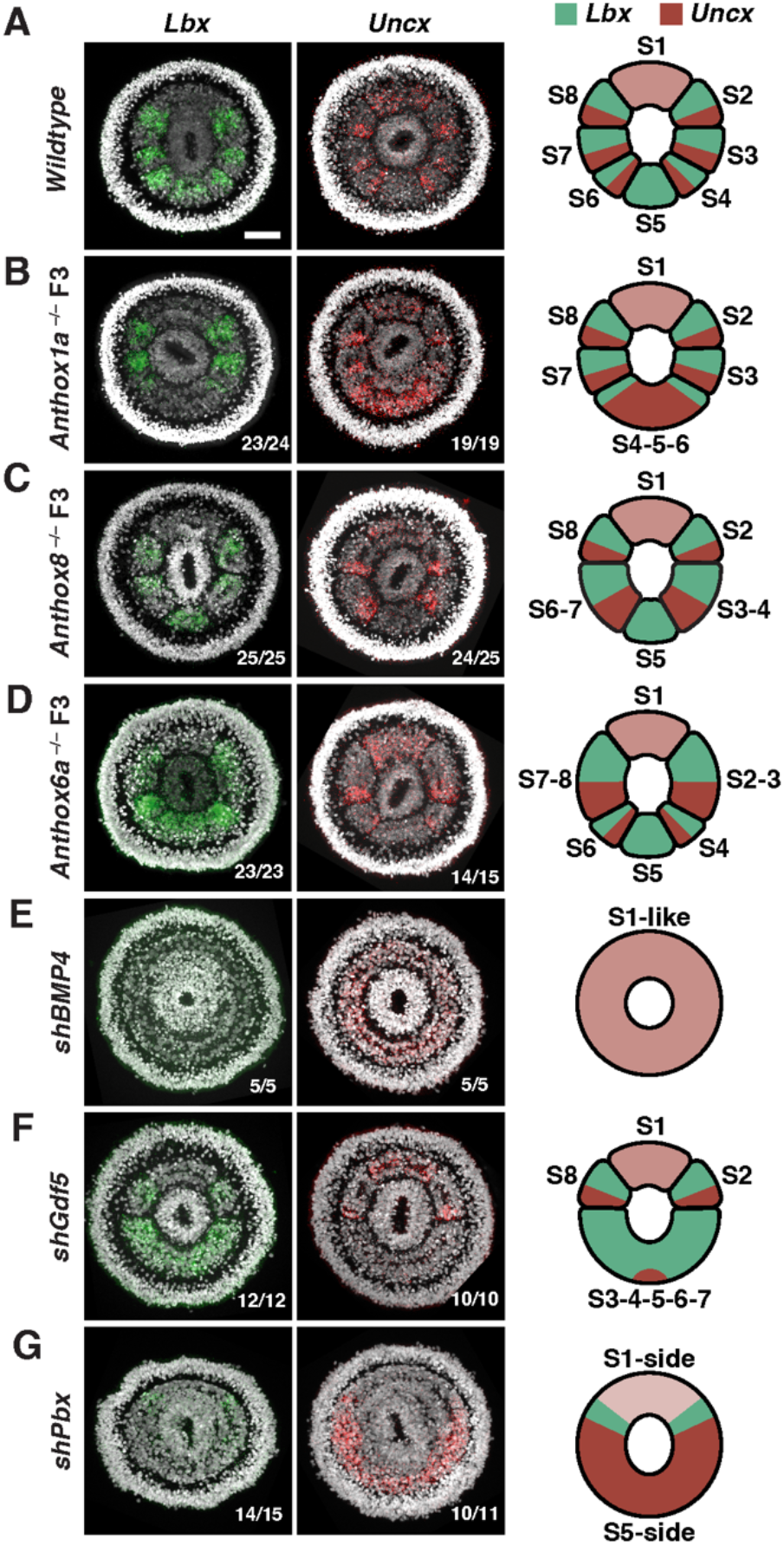
The *Nematostella* segment polarity program is under the cooperative regulation of BMP signal and Hox-Gbx genes. (**A** to **D**) *Lbx-Uncx* expression patterns under *wildtype* (**A**), *Anthox1a*^-/-^ (**B**), *Anthox8*^-/-^ (**C**) and *Anthox6a*^-/-^ (**D**) genetic backgrounds. (**E** to **G**) *Lbx-Uncx* expression patterns in animals injected with *shBMP4* (**E**), *shGdf5* (**F**) and *shPbx* (**G**). Embryos were collected at 72hpf, and the oral views were displayed. Cartoon diagrams on the right depicts the polarity patterns observed and illustrate the segment names. Numbers on the lower right corner indicate the observed expression patterns in the total number of animals imaged. Scale bar, 50μm.

Importantly, Hox genes are homogenously expressed within developing endomesodermal segments and their activities are thus unlikely to be sufficient to generate polarized *Lbx*-*Uncx* stripes. In *Nematostella*, BMP signaling forms an activity gradient across the endomesoderm which directly activates Hox-Gbx genes and defines their expression territories (Genikhovich et al., 2015; Kraus et al., 2016; Leclere and Rentzsch, 2014; Rentzsch et al., 2006; Wijesena et al., 2017). We therefore suspected the existence of an additional regulatory input from the BMP pathway upstream of *Lbx-Uncx* polarity. This hypothesis was further strengthened by the existence of a phosphorylated Smad1/5 (pSmad1/5) binding peak upstream of the *Lbx* locus from ChIP-seq data (Knabl et al., 2022), indicating that *Lbx* is a direct target of the BMP pathway. To test this experimentally, we first abolished BMP activity by knocking down major BMP pathway components in *Nematostella* including *Bmp4*, *Bmp5*, *Chordin* and *Smad1/5* (**Fig. 5E**, **Fig. S5A** to **C**). In line with previous work, no endomesodermal segments were formed in shRNA-injected larvae (**Figure S5A**). The resulting unsegmented endomesoderm lacked *Lbx* expression, and uniformly expressed *Uncx* (**Fig. 5E**, **Fig. S5B** and **C**). Furthermore, the segment S1 marker *Arp6* expanded across the endomesoderm, suggesting that the entire tissue was transformed into an S1-like ground state in the absence of BMP activity (**Figure S5H**). Next, by knocking down the TGF-β pathway signaling molecule *Gdf5*, we examined the effect of a weakened BMP gradient on segment polarity. In the absence of *Gdf5*, the flattened BMP activity gradient failed to activate the high-threshold Hox genes *Anthox1a* and *Anthox8*, while the low-threshold genes *Anthox6a* and *Gbx* were unperturbed, resulting in the fusion of segments S3-S7 (**Fig. 5F**, (Knabl *et al*., 2022)). Contrary to the fused segments observed in different Hox mutants, these S3-S7 fusions displayed an altered polarity, where *Lbx* became ubiquitous and *Uncx* was undetectable (**Fig. 5F)**. Collectively, these results support a model in which a stepwise decreasing activation signal from Hox-Gbx genes counteracts the continuously decreasing repressive signal from the BMP pathway to generate segmentally polarized *Lbx* stripes along the directive axis (**Fig. S5I** to **K**). *Lbx* is thus only expressed at the S1-sided borders of each Hox-Gbx expression domain, where repressive BMP activity is lowest and thus overcome by the activation input.

To further validate the model, we examined the expression of polarity markers in larvae injected with two independent shRNAs targeting the Hox binding partner *Pbx* (**Fig. 5G**, **Fig. S5D** to **G**). Knockdown of *Pbx* disrupted the function of all Hox-Gbx genes acting downstream of the BMP pathway, leading to the formation of an unsegmented endomesoderm at the planula stage (**Fig. S5D**). Although morphologically resembling the *Bmp4* KD condition (**Fig. S5A**), *Pbx* KD larvae exhibited a distinct transcriptomic profile (**Fig. S5E** to **G, Table S3**). *Anthox8*, which requires Pbx-dependent self-activation, was significantly downregulated, whereas *Anthox6a* and *Gbx* were upregulated (**Fig. S5G**; He *et al*., 2018). Moreover, despite the lack of segment boundaries, *Arp6* expression was still restricted to a polar region corresponding to segment S1, suggesting that a BMP activity gradient that represses the ground state identity remained functional in the absence of *Pbx* (**Fig. S5H**). Interestingly, the unsegmented endomesoderm in *Pbx* KD larvae still displayed polarized *Lbx-Uncx* patterns: two *Lbx* stripes flanking the *Arp6* domain were observed, restricting a high level of Uncx expression to the S5-sided endomesoderm (**Fig. 5G**). These results indicate that *Gbx* likely retains some gene regulatory activity in the absence of *Pbx*, resulting in the establishment of a modified *Lbx-Uncx* polarity in the unsegmented endomesoderm (**Fig. S5L**).

### *Lbx* controls polarized positioning of retractor muscles in *Nematostella*

To investigate the developmental requirements for segment polarization, we next generated two mutant alleles that disrupt the major functional domains of the LBX protein (**Figure 6A, S7B and C**). *Lbx* homozygous mutants did not display segmentation defects at 72hpf and metamorphosed into polyps with four properly positioned tentacle primordia (**Figure S6A** and **B**). However, these mutant polyps exhibited abnormal elongation of the oral-aboral axis, possessing a shortened body column and stubby tentacles (**Figure S6C** to **F**). Consistent with these defects, *Lbx* homozygous polyps were unable to feed effectively and were quickly outcompeted by *wild-type* and *Lbx*/+ siblings in mixed cultures (**Figure 6B**).

**Figure 6.**
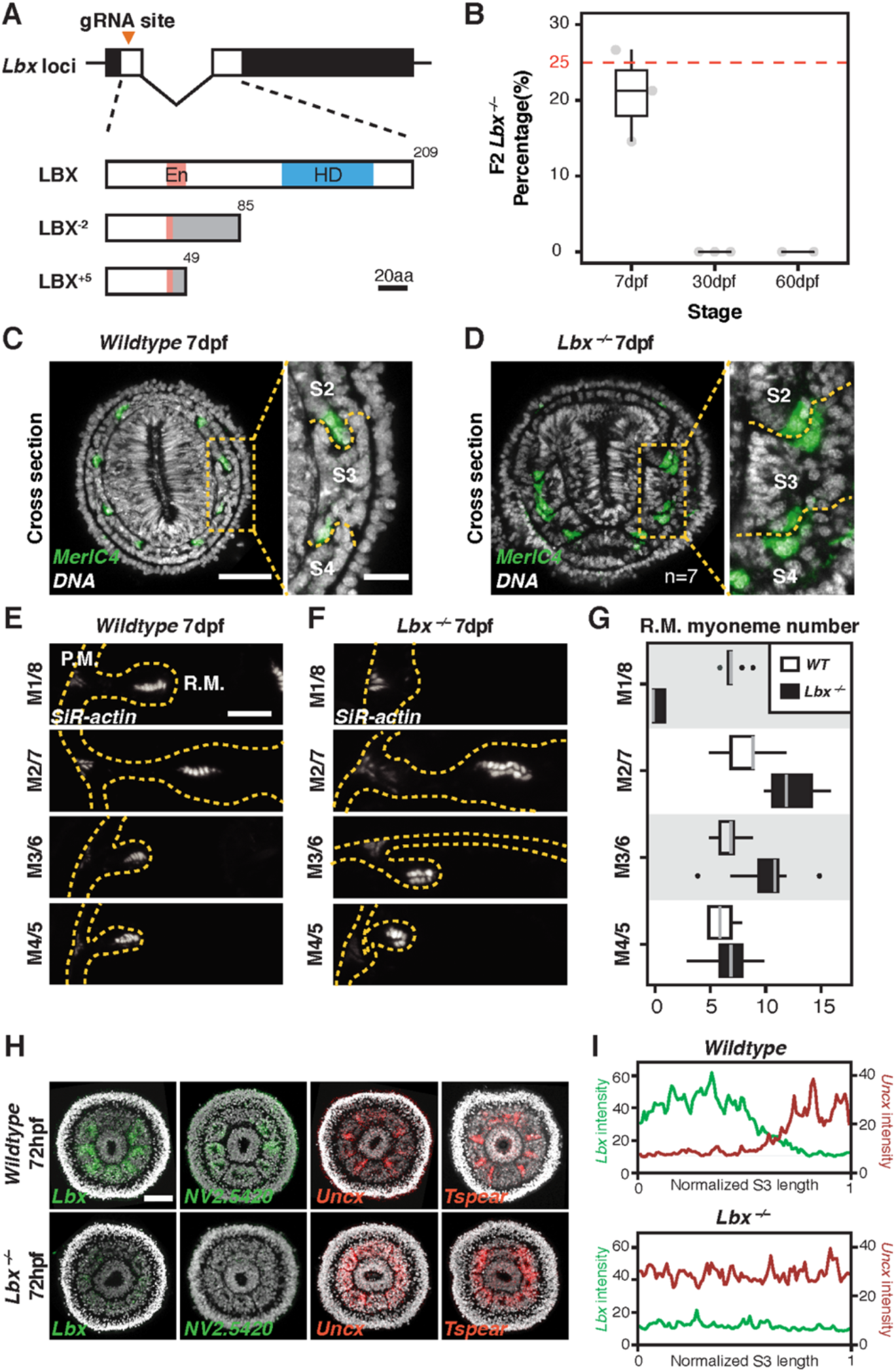
*Lbx* Loss of function resulted in disrupted segment polarity. (**A**) A single guide RNA was selected for CRISPR mutagenesis to generate truncated forms of LBX in two different mutant alleles. (**B**) Quantification of genotypes at 7dpf, 30dpf and 60dpf from F1 heterozygous crosses. (**C** and **D**) Cross section view of *wildtype* (C) and *Lbx* mutant (D) polyps at 7dpf. Retractor muscle cells were labeled with probes against *myosin essential light chain 4* (*Melc4*). Zoomed in images showing retractor muscles at the junctions between segment S2, S3 and S4. Scale bars, 50μm for cross section, 15μm for zoomed in images. (**E** and **F**) F-actin staining showing myoneme arrangement of retractor muscles residing in different mesenteries between *wildtype* (E) and *Lbx* mutant (F) polyps at 7dpf. Scale bar, 10μm. (**G**) Quantification of retractor muscle myoneme numbers between *Lbx* mutant and *wildtype* polyps. (**H**) Expression of segment polarity genes in *wildtype* and *Lbx* mutant planula larvae at 72hpf. Scale bar, 50μm. (**I**) Quantification of *Lbx-Uncx* FISH signal along the normalized length of segment S3 in *wildtype* and *Lbx* mutant planula larvae at 72hpf.

The observations above suggest that *Lbx* mutants may have additional defects and prompted us to investigate other segmentally polarized features of developing polyps, including the positioning of the retractor muscles (RMs). In *wild-type* animals, RMs formed on the S1-side of mesentery pairs m2/m7, m3/m6 and m4/m5 and on the S5-side of mesenteries m1/m8, indicated by the expression of the marker gene *MelC4* (**Figure 6C**). In *Lbx* mutant polyps, however, each mesentery displayed symmetric RM patterning. Mesentery pair m1/m8 no longer formed RMs, while the rest of the mesenteries formed duplicated RMs, specifically on their S5-side (**Figure 6D**). At the morphological level, the duplicated RMs exhibited inverted polarity, with their actin-rich basal myonemes facing each other, separated by the mesoglea (**Figure E** to **G**). Since each mesentery is formed by joining two compartments from neighboring segments, the RM patterning defects we observed in *Lbx* mutants are consistent with aberrant segment polarity. Indeed, the expression of *Lbx*, *Nv2.5420* and other S1-sided genes were diminished in *Lbx* homozygous mutants (**Figure 6H**). Conversely, *Uncx* became ubiquitously expressed across the endomesoderm, together with other S5-sided genes such as *Tspear* and *Thsd4*, suggesting that normal segment polarity collapsed in the absence of *Lbx* (**Figure 6H** and **I**).

### An Evolutionarily conserved segmental polarity program in *Nematostella*

The identification of the *Lbx-Uncx* segment polarity module in a cnidarian animal prompted us to further investigate the evolutionary origin of both genes. By performing reciprocal BLAST searches using the full-length as well as homeobox sequences of *Nematostella* LBX and UNCX proteins, we confirmed the presence of both genes in the genomes of five additional anthozoan species, including *Montipora capitata*, *Acropora digitifera*, *Stylophora pistillata*, *Xenia spp*. and *Scolanthus callimorphus* (**Figure S7C** and **D**, **Table S3**). To our surprise, non-anthozoan classes including Scyphozoa (represented by *Aurelia aurita* and *Nemopilema nomurai*), Cubozoa (represented by *Tripedalia cystophora* and *Morbakka virulenta*) and Hydrozoa (represented by *Hydra magnipapillata*, *Hydra viridissima*, *Hydractinia symbiolongicarpus* and *Hydractinia echinata*) do not possess *bona fide Lbx* or *Uncx* (**Figure 7A**, **Table S3**). Furthermore, we were unable to identify definitive *Lbx-Uncx* homologs in the basal metazoan phyla Porifera (represented by *Ephydatia muelleri* and *Amphimedon queenslandica*), Ctenophora (represented by *Hormiphora californensis*, *Pleurobrachia bachei* and *Mnemiopsis leidyi*) and Placozoa (represented by *Trichoplax adhaerens*) (**Table S3**). In contrast, phylogenetic reconstruction using full length amino acid sequences strongly supports the homology between cnidarian LBX-UNCX proteins and their bilaterian counterparts (**Figure S7A and B**). Taken together, these results indicate that *Lbx* and *Uncx* first emerged in the Cnidaria-Bilateria common ancestor, likely through a gene duplication event, and were partially or completely lost in several unsegmented lineages, including non-anthozoan cnidarians, acoels, nematodes as well as bryozoans (**Figure 7A, Table S3**). Interestingly, both *Lbx* and *Uncx* have been implicated to function as polarity genes in segmented bilaterians including arthropods, annelids and vertebrates (De Graeve et al., 2004; Dray et al., 2010; Jagla et al., 1997; Mansouri et al., 2000; Saudemont et al., 2008; Treffkorn et al., 2018). Based on these observations, it is plausible that an Lbx/Uncx-like segment polarity module existed in the cnidarian-bilaterian common ancestor, which was subsequently lost or modified in diverse animal lineages during body plan evolution.

**Figure 7.**
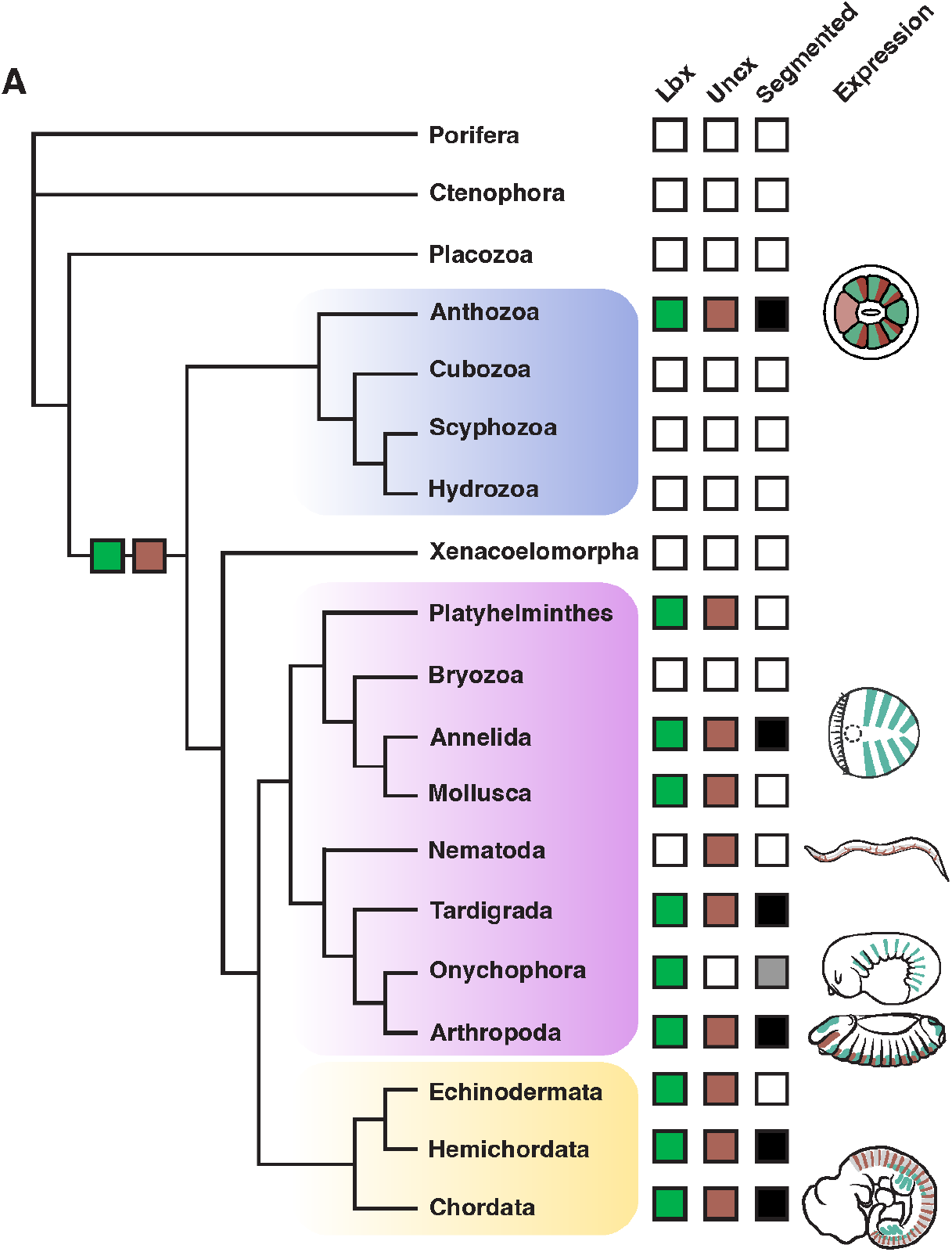
Evolution of the segmental polarity program. Phylogenic tree depicting the evolutionary relationships between major metazoan phyla and the presence or absence of segmented body plans and *Lbx-Uncx* genes in the respected genomes. The *Lbx-Uncx* expression patterns of representative species were adapted from previous publications.

## Discussion

### *Nematostella* segment identity is determined via a Hox-Gbx hierarchy

We previously proposed that the *Nematostella* Hox-Gbx hierarchy generates a binarized identity outcome for each segment: either tentacle forming or non-tentacle forming (**Figure 3A**). However, this simplified conception of segment identity fails to explain other segment-specific features, such as the development thickened primary mesenteries which only form at the boundaries between segments S2 and S3 and S7 and S8. In addition, as the primary polyp grows, new tentacles emerge following a stereotypic pattern, and non-tentacle bearing segments will eventually permit tentacle growth once the polyp reaches a certain size (Ikmi *et al*., 2020). The development of Endo-atlas now permits a deeper understanding of *Nematostella* segment identity at the molecular level. Polar segment S1 possesses a unique transcriptomic profile, being the only segment that does not express any Hox-Gbx genes, thus representing the “ground state” of endomesoderm (**Figure S5I**). Segment S1 is also highly neurogenic and has been shown as the first segment to differentiate *GLWamide*^+^ neurons (Watanabe *et al*., 2014). The even-numbered segment pairs S2/S8 and S4/S6, being tentacle forming segments, share somewhat similar transcriptomic profiles, exemplified by the co-expression of *Dmbx2*, *Zic3*, *Col6A5* and *Nv2.2940* (**Figure 3C**). However, each of these segment pairs also possess some distinguishing markers. Segments S2 and S8 co-express the homeobox-containing genes *Dmbx1* and *Q50*, while segments S4 and S6 co-express *Anthox7*, *TgfR3*, *F5* and *Cdx* (**Figure 3D**, **Figure S3A**). The remaining non-tentacle bearing segments S3/S7 and S5 also exhibit drastically different transcriptomic profiles, with the polar segment S5 possessing many unique segment markers: *Efemp1*, *F8* and *Nv2.7863* among others (**Figure 3E**, **Figure S3A**). Consequently, each segment possesses a unique molecular identity along the directive axis. In general, the even-numbered segments are more similar to each other, while the odd-numbered segment pairs show fewer similarities. These observations suggest that *Nematostella* Hox-Gbx genes share some common targets but also possess unique sets of downstream targets, reminiscent of their bilaterian counterparts.

### Segment polarity in *Nematostella* is cooperatively determined by BMP and Hox-Gbx genes

As demonstrated by previous studies, BMP signaling is instrumental during directive axis establishment and endomesoderm patterning in *Nematostella* (Leclere and Rentzsch, 2014; Rentzsch *et al*., 2006; Wijesena *et al*., 2017). Prior to segment formation, BMP signaling forms an activity gradient across the endomesoderm, which peaks at the future segment S5 and gradually diminishes towards segment S1 (Genikhovich *et al*., 2015). Hox-Gbx genes are activated along this gradient at different thresholds. The expression boundaries of each Hox-Gbx genes then determine the locations of physical boundaries, resulting in the formation of 3 segment pairs (S2/S8, S3/S7 and S4/S6) as well as two polar segments (S1 and S5) along the directive axis (He *et al*., 2018). In the current work, we further demonstrate the existence of a paraxial molecular polarity consisting of two homeobox containing genes: *Lbx* and *Uncx* (**Fig. 4**). These factors are expressed in complementary expression territories within each segment pair and convey distinct developmental potentials (**Fig. 4**, **Fig. 6**).

How are subsegmental *Lbx* and *Ubx* expression domains established? We propose that *Lbx* receives counteracting regulatory inputs from the BMP pathway and Hox-Gbx genes (**Fig. S5I**). In this model, BMP signaling represses *Lbx* expression with decreasing strength along its activity gradient, whereas Hox-Gbx genes are able to activate *Lbx* expression with different strengths. In this scenario, BMP’s capacity to repress Lbx is lowest at the S1-side of each segment. In parallel, we propose that the Hox-Gbx activation strength decreases along the same axis, with *Gbx* having the weakest capacity to induce *Lbx* near the S1-side. Consequently, within each segment pair, *Lbx* is only expressed in the S1-sided regions where the decreasing BMP activity can be overcome by the activation input provided by Hox-Gbx genes. Further, polar segment S1, which lacks Hox-Gbx expression, does not turn on *Lbx* during normal development, and displays the same “ground state” molecular profile (*Arp6*^+^, *Uncx*^+^) as the entire unsegmented endomesoderm under BMP KD conditions (**Fig. 5**, **Fig. S5H**).

Contrary to expectations, we found that knocking down the Hox binding partner *Pbx* does not recapitulate BMP KD conditions at the molecular level (Knabl *et al*., 2022). In fact, despite the lack of all physical boundaries, segment S1 identity, as marked by *Arp6* expression, is still confined to the S1-sided polar region, and the paraxial segment polarity program remains active in *shPbx* injected larvae (**Fig. 5G**, **Fig. S5D** and **H**). These observations suggest that the axial patterning activity of BMP signaling is not entirely dependent on the proper functioning of Hox-Gbx genes. Moreover, *Gbx* appears to retain its activation input into the *Lbx* locus in the absence of *Pbx*, resulting in the formation of two weak *Lbx* stripes flanking the unsegmented S1 territory (**Fig. S5L**). Combined, these observations support the idea that segment polarity establishment in *Nematostella* is the outcome of combinatorial regulatory interactions between an upstream signaling gradient (BMP) and its direct downstream compartment identity genes (Hox-Gbx). Together, they determine polarity gene expression domains prior to and independent of the establishment of physical segment boundaries.

### The evolution of the *Lbx-Uncx* segment polarity module

*Lbx* and *Uncx* are both conserved homeobox-containing transcription factors and are known to exhibit segmentally polarized expression patterns during embryogenesis in various bilaterian lineages. Originally discovered in *Drosophila* and classified as a segment polarity gene, *Lbx* encodes a NK class homeobox protein that specifies muscle identity during body segmentation in arthropods and vertebrates (De Graeve *et al*., 2004; Jagla et al., 1995; Juarez-Morales et al., 2021; Ochi and Westerfield, 2009; Wotton et al., 2008). In the annelid *Platynereis*, *Lbx* demarcates future posterior segment boundaries and forms complementary stripes with other polarity genes including *Engrailed*, *Wnt1* and *Tlx* (Saudemont *et al*., 2008). This process is dependent on the upstream Hedgehog signal and occurs prior to the formation of physical segment boundaries (Dray *et al*., 2010). A similar *Lbx* expression pattern was reported in Onychophora (velvet worms), where *Lbx* stripes are detected anterior to the growing segmental furrows (Treffkorn *et al*., 2018). Chordates, however, have lost segmentally polarized *Lbx* expression and instead rely on the gene to specify migratory hypobranchial and appendicular muscle progenitors (Kusakabe et al., 2020; Ochi and Westerfield, 2009). This transition of the *Lbx* expression pattern coincides with the emergence of a different mode of body segmentation in chordates, where somatic musculature does not develop in a segmentally polarized manner. *Uncx* is a member of the PRD class homeobox genes and was initially identified in C. elegans as *Unc-4*. In protostomes, *Uncx* function is largely restricted to the nervous system, where it specifies motor neuron identity, sometimes in a repeated, segmentally polarized fashion (Cho and Park, 2008; Lacin et al., 2020; Miller et al., 1992; Walthall, 1995). In contrast, the mammalian *Unc-4* homolog *Uncx4.1* serves as one of the best characterized posterior identity markers of the developing somites, where it instructs the formation of distal ribs, transverse processes and pedicles of the neural arches on vertebrae (Leitges et al., 2000; Mansouri *et al*., 2000; Neidhardt et al., 1997; Schragle et al., 2004).

Given that *bona fide* Lbx and Uncx homologs cannot be identified in basal metazoan clades including Ctenophora, Porifera and Placozoa, it is likely that both genes first evolved in the Cnidaria-Bilateria common ancestor, in concert with the emergence of a bi-radially segmented body plan (**Fig. S7**, **Fig. 7**, **Table S3**). However, despite being implicated as segment polarity genes in diverse bilaterian systems, *Lbx* and *Uncx* have not been shown to function together during polarity establishment in animals other than *Nematostella*. One potential explanation is that there has been convergent evolution in the patterning of somatic musculature and nervous systems in diverse metameric animal lineages. Consequently, important cell identity regulators such as *Lbx* and *Uncx* could be frequently employed in a segmentally polarized manner and thus exhibit similar expression patterns under the control of distinct upstream signals. Alternatively, we postulate that the *Lbx-Uncx* pair could represent an ancient and rudimentary segment polarity module that has been modified and rewired under the control of different signaling pathways to pattern the incredible diversity of metameric structures that have arisen during bilaterian evolution.

## ACKNOWLEDGMENTS

This work was performed to fulfill, in part, requirements for S.H.’s thesis research in the Graduate School of the Stowers Institute. All authors listed have read and approved the manuscript. The authors would like to thank Stowers Institute Aquatics team for assistance with animal husbandry, Sequencing and Discovery Genomics team for library preparation and sequencing, all Gibson Lab members for helpful discussions and Prof. Robb Krumlauf (Stowers Institute), Prof. Hopi Hoekstra (Harvard University) for comments and suggestions regarding the work.

## Funding

S.H., S.C. and M.C.G. are funded by the Stowers Institute for Medical Research. W.S. and T.W. are funded by National Institutes of Health grants R01HG007175, U24ES026699, U01CA200060, U01HG009391, and U41HG010972.

## Author Contributions

Conceptualization: S.H. and M.C.G. Experimentation: S.H. Analysis: W.S. and S.C. Computational resources: T.W. Writing: S.H. and M.C.G. All authors provided feedback on the manuscript.

## Competing interests

The authors declare no competing interest.

## Data and materials availability

Original data underlying this manuscript can be downloaded from the Stowers Institute for Medical Research Original Data Repository: http://www.stowers.org/research/publications/libpb-1623. All NGS sequencing results are available at GEO: https://www.ncbi.nlm.nih.gov/geo/query/acc.cgi?acc=GSE173786. The *Nematostella* Endo-atlas can be accessed at: http://endoatlastest-env.eba-qtpfn7qz.us-east-1.elasticbeanstalk.com/. The NV2 genome and transcriptome used in this study are hosted on SIMRbase: https://genomes.stowers.org/.

## SUPPLEMENTARY MATERIALS

STAR Methods

Figure S1–7

Tables S1, S2 and S3

**Figure S1.**
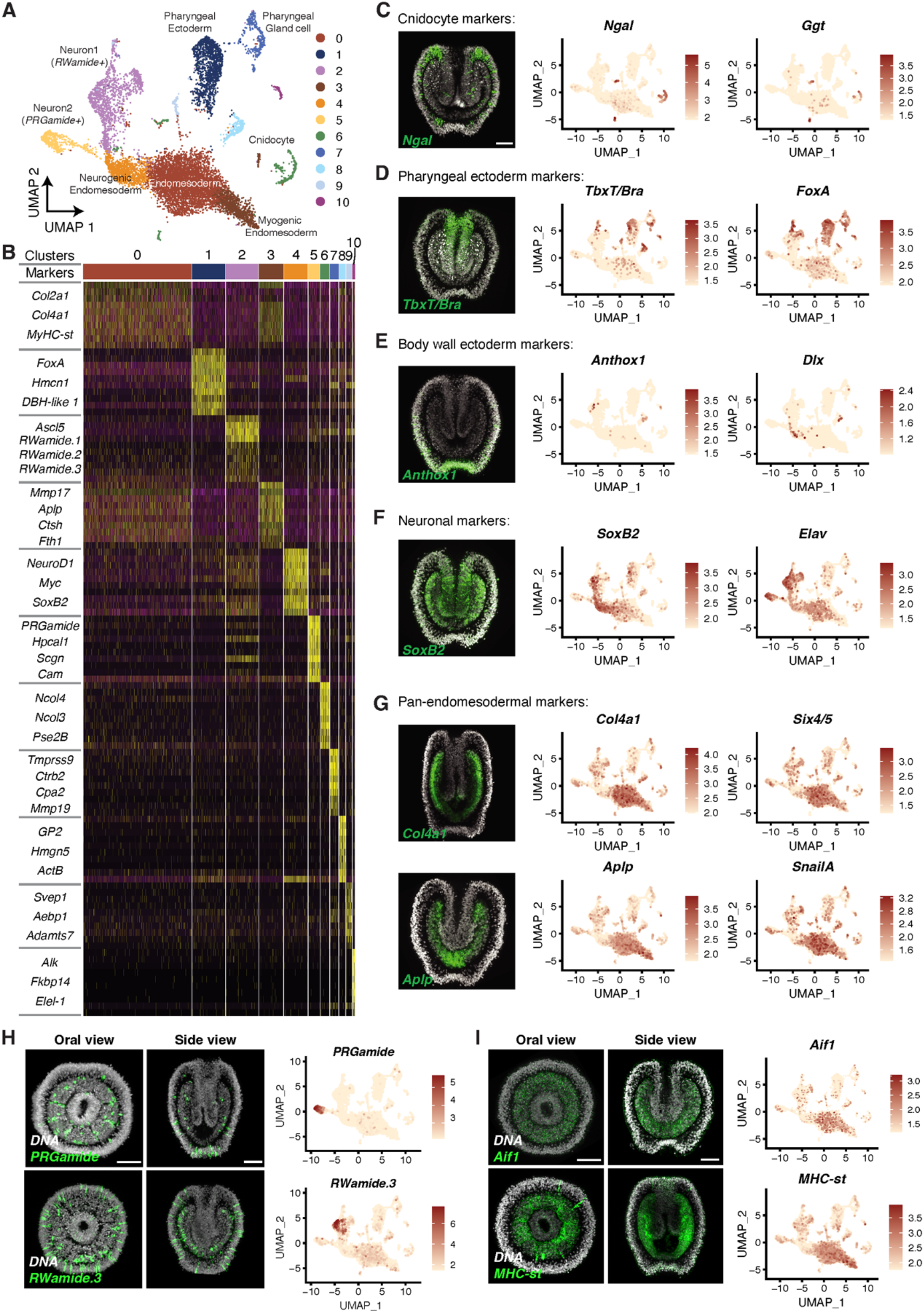
Annotating cell clusters using known and novel marker genes, related to Figure 1. (**A**) UMAP projection showing all cell clusters identified from scRNA-seq of the endomesoderm isolates. (**B**) Top 10 markers for each cluster, identified by Seurat. Gene names of a subset of markers was listed on the left. (**C** to **G**) Fluorescent *in situ* hybridization and single cell feature plots confirming the expression patterns of different cluster markers. (**C**) Non-pharyngeal ectodermal markers such as *Anthox1* and *Dlx* were barely detected in the dataset, indicating the removal of the majority of ectodermal tissue through the isolation process. Scale bar, 50μm. (**H** and **I**) Fluorescent *in situ* hybridization and single cell feature plots confirming the expression patterns of different cell type markers in the endomesoderm. Scale bars, 50μm.

**Figure S2.**
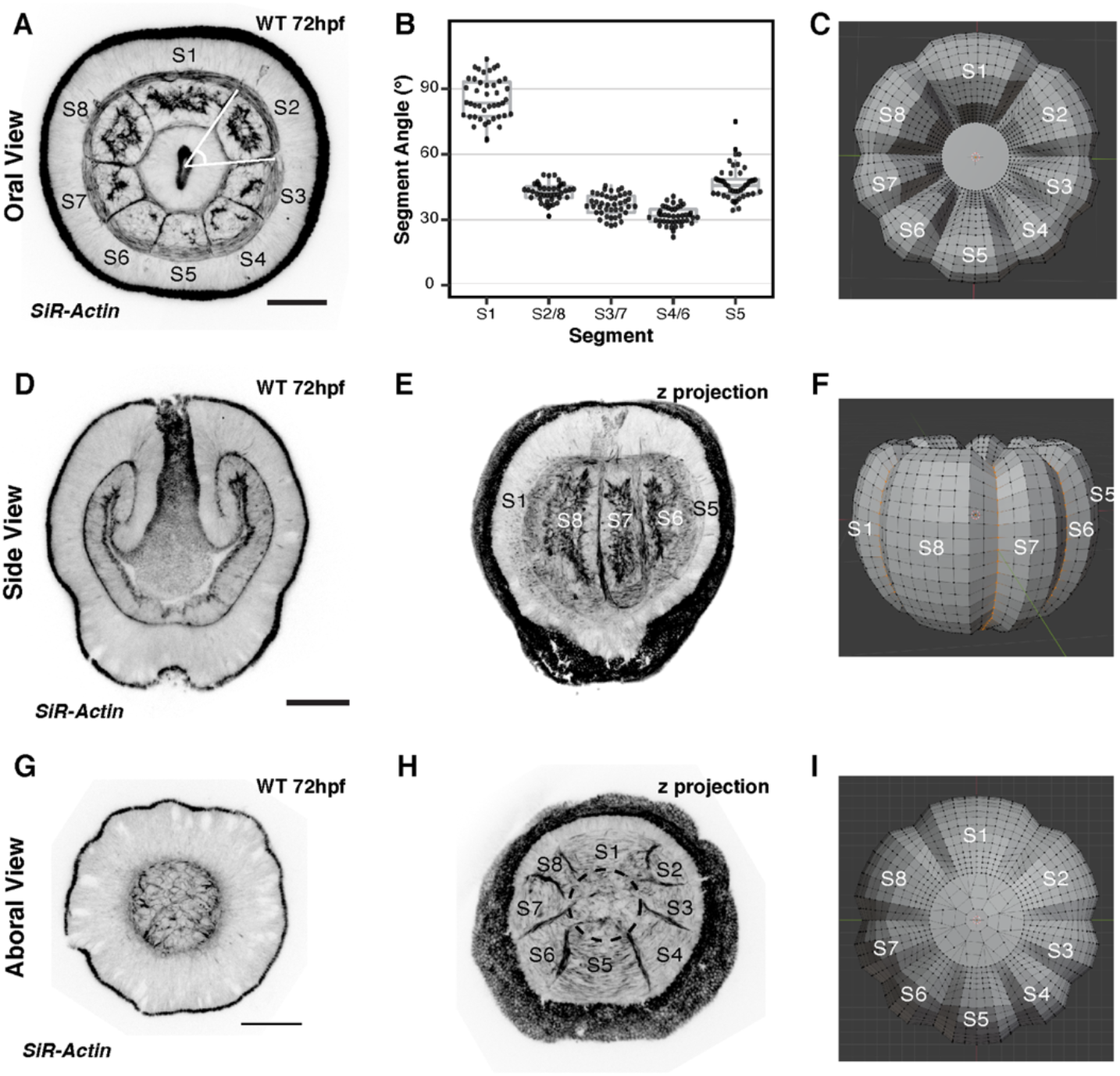
Generating a 3-D model of *Nematostella* endomesoderm based on the morphology and geometry of the tissue, related to Figure 2. (**A** to **C**) Oral view comparison of the actual embryo and the model. Size of each segment in the model was determined based on the angle measurements in real larvae. (**D** to **F**) Side view comparison of the actual embryo and the model. Most of the endomesoderm as is completely segmented at this time point as segment boundaries are visible along the oral-aboral axis. (**G** to **I**) Aboral view comparison of the actual embryo and the model. The aboral most endomesoderm remains unsegmented at 72hpf, which is also depicted in the model. Scale bars, 50μm.

**Figure S3.**
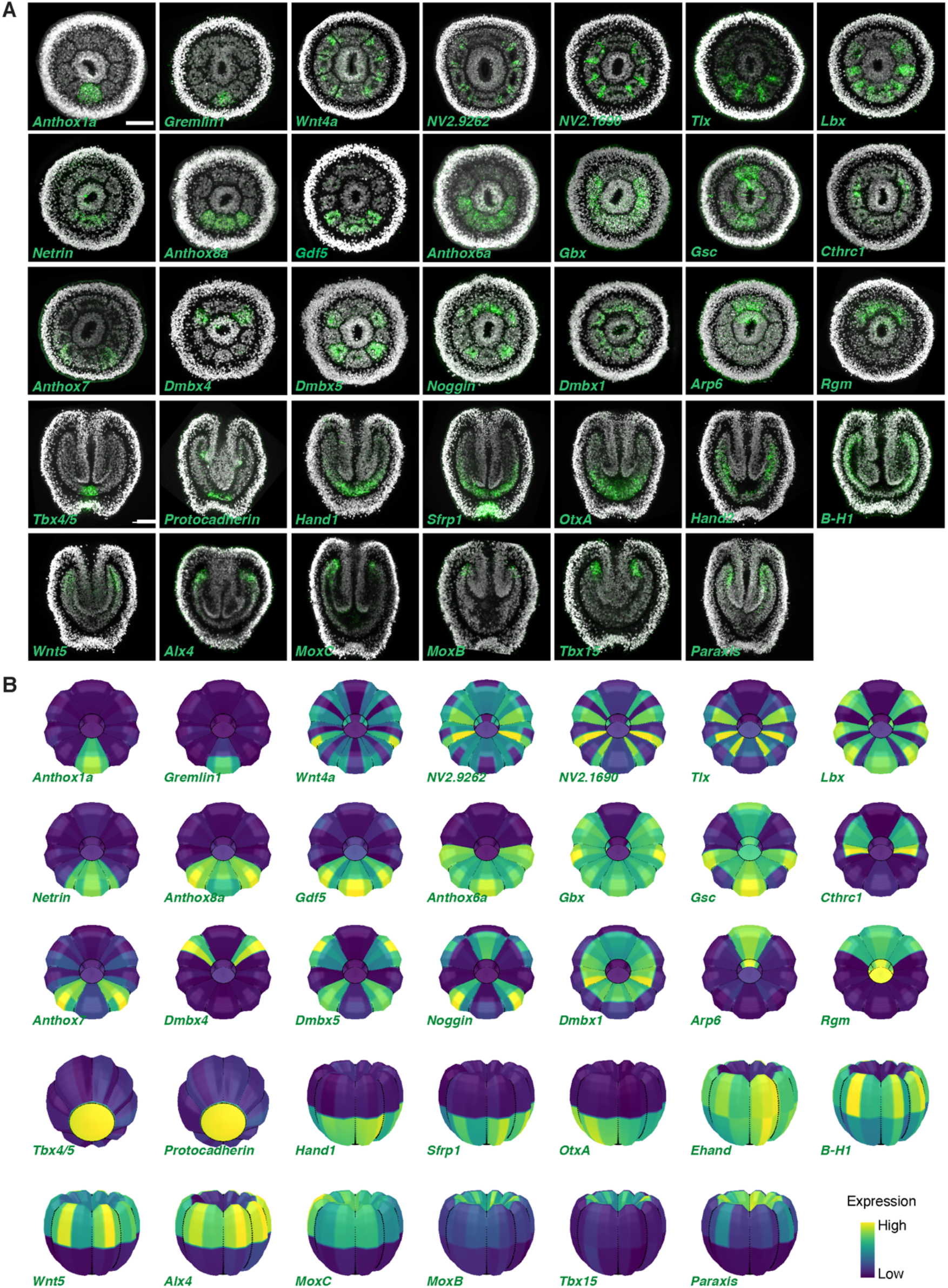
Expression patterns and Endo-atlas predictions of all 34 landmark genes used in this study, related to Figure 2. (**A**) FISH results of landmark genes with distinct patterns in the developing endomesoderm. Scale bars, 50μm. (**B**) Endo-atlas predictions of the same 34 genes. Color scale indicates the relative expression value (yellow, high; blue, low).

**Figure S4.**
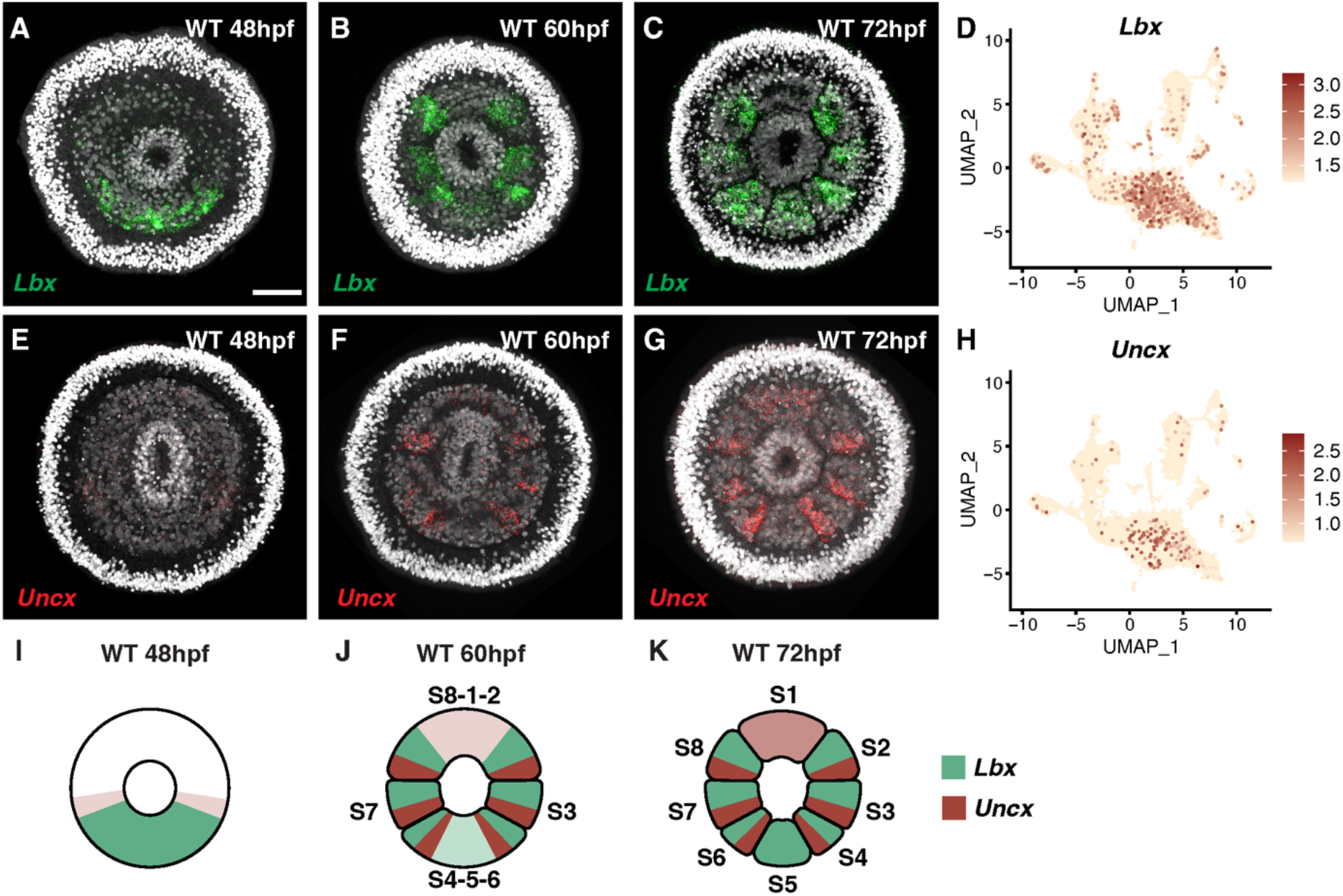
The dynamic expression of *Lbx-Uncx* during segment formation, related to Figure 4. (**A** to **C**) *Lbx* expression in 48hpf, 60hpf and 72hpf planula larvae. Scale bar, 50 μm. (**D**) Expression of *Lbx* in the single cell dataset at 72hpf. (**E** to **G**) *Uncx* expression in 48hpf, 60hpf and 72hpf planula larvae. (**H**) Expression of *Uncx* in the single cell dataset at 72hpf. (**I** to **K**) Cartoon illustration of the temporal and spatial expression of *Lbx-Uncx*. As shown in (**B** and **F**), subsegmental *Lbx* stripes flanking the future segment S1 appear prior to the formation of physical segment boundaries between S1/S2 and S1/S8, whereas *Uncx* stripes flanking segment S5 appear prior to the formation of segment S5.

**Figure S5.**
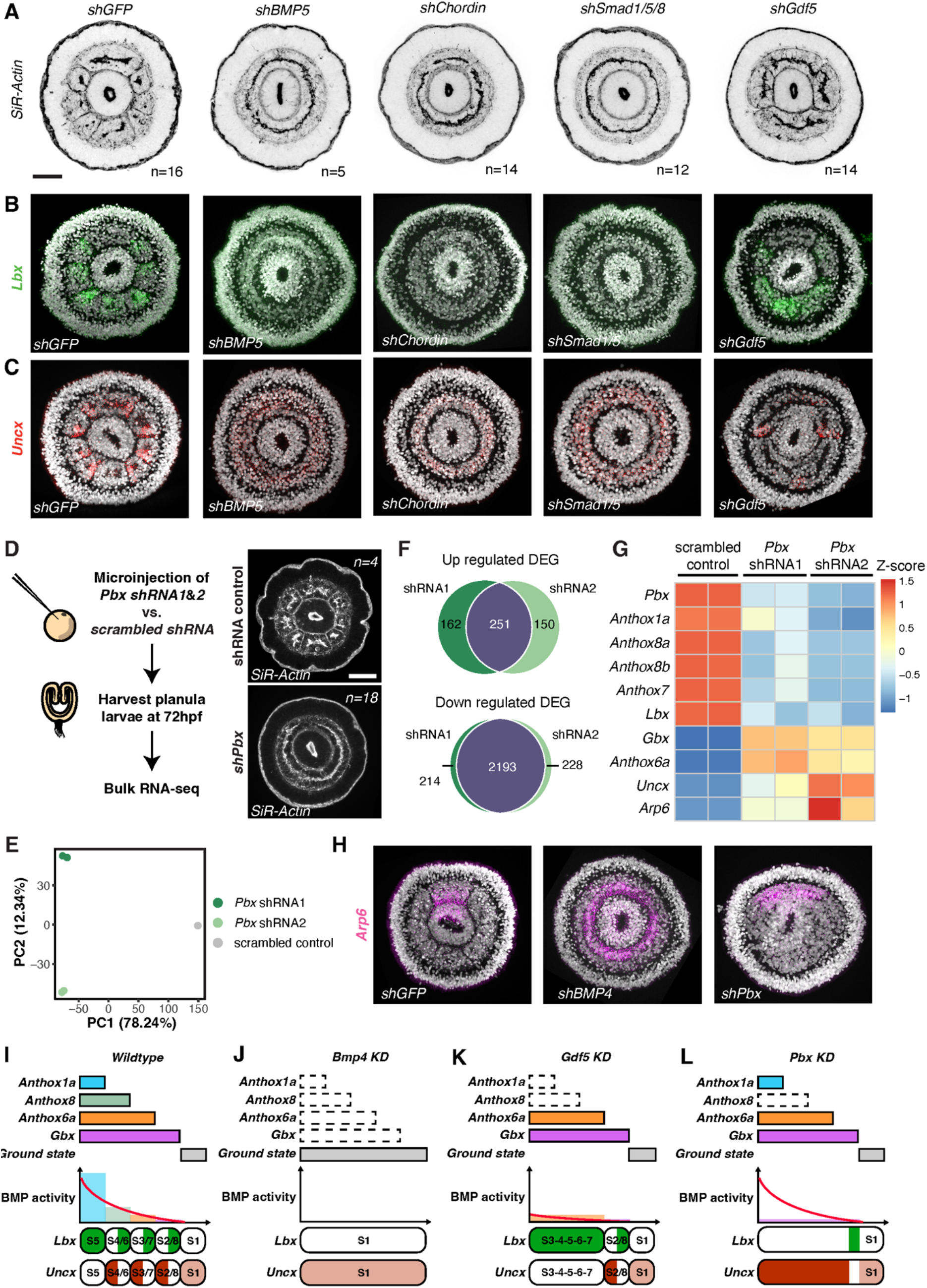
The upstream regulation of *Lbx-Uncx* polarity, related to Figure 5. (**A**) The oral view of *Nematostella* larvae 72 hours after injection of *shGFP*, *shBMP5*, *shChordin*, *shSmad1/5* and *shGdf5*. Scale bar, 50μm. (**B** and **C**) *Lbx-Uncx* expression in animals injected with different shRNAs. (**D**) Experimental design to evaluate the global transcriptomic changes induced by *Pbx* knockdown. Two independent shRNAs targeting *Pbx* were used. Side panels show F-actin staining of 72hpf planulae injected with scrambled control shRNA and *shPbx*, respectively. Scale bar, 50μm. (**E**) Principal Component Analysis of the RNA-seq results. (**F**) Venn diagrams comparing significantly up and down regulated genes for each shRNA targeting *Pbx*. (**G**) Heatmap of Z-score of homeobox containing genes under different shRNA injection groups. *Lbx* is down regulated after *Pbx* knockdown while *Uncx* is up regulated. (**H**) FISH results showing the expression of segment S1 marker *Arp6* in control, *shBMP4* and *shPbx* injected planulae. Scale bars, 50μm. (**I** to **L**) Cartoon illustrations of the regulatory logic upstream of the segment polarity program in *wildtype*, *BMP4* KD, *Gdf5* KD and *Pbx* KD conditions.

**Figure S6.**
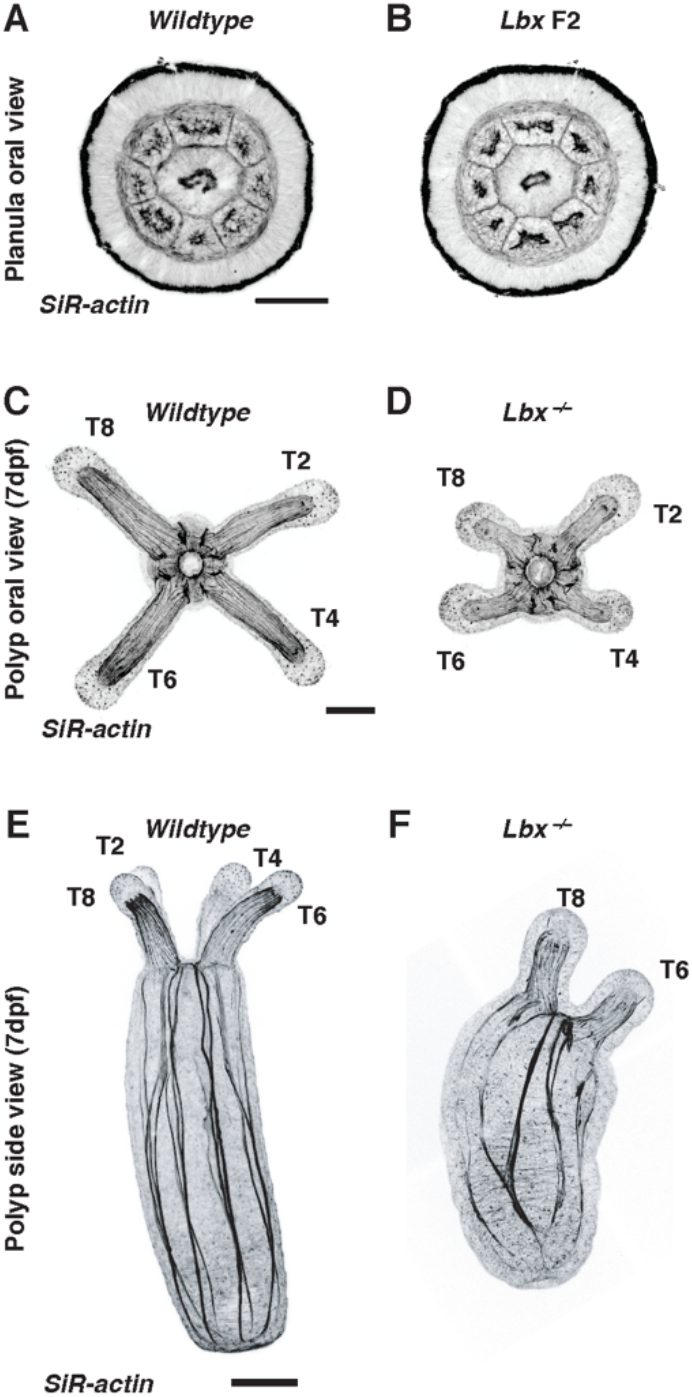
Morphological analysis of *Lbx*^-/-^ mutants, related to Figure 6. (**A-B**) Planula stage morphological comparison between *Lbx* homozygous mutants and their *wildtype* siblings. Scale bars, 50μm. (**C-D**) Polyp stage morphological comparison between *Lbx* homozygous mutants and their *wildtype* siblings. Scale bars, 100μm.

**Figure S7.**
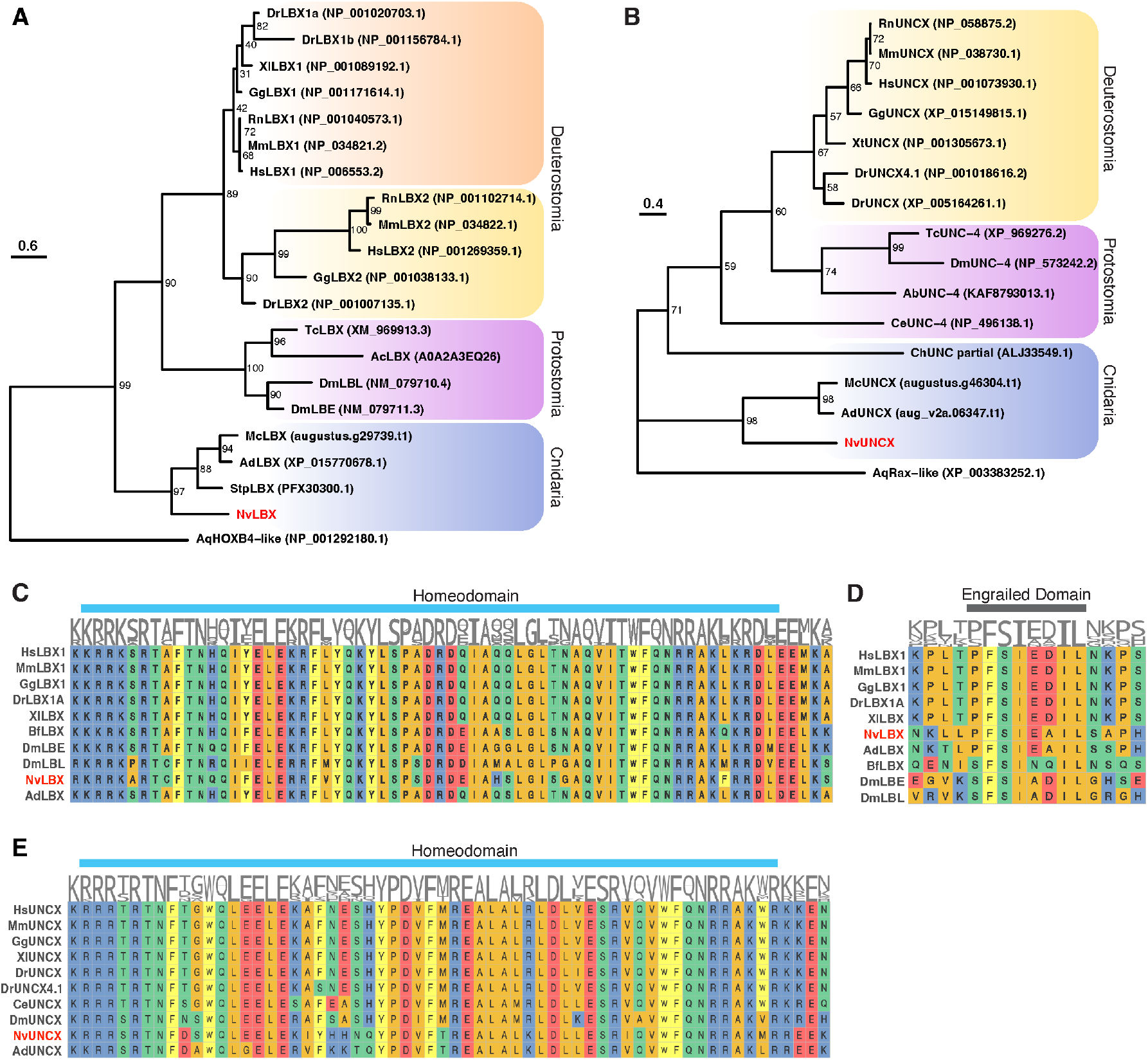
*Nematostella* LBX and UNCX are conserved homeodomain-containing proteins, related to Figure 7. (**A**) Maximum-likelihood phylogenic tree of LBX protein. (**B**) Maximum-likelihood phylogenic tree of UNCX protein. (**C**) Sequence alignment of the homeodomain of LBX proteins from different species. (**D**) Sequence alignment of the engrailed domain of LBX proteins from different species. (**E**) Sequence alignment of the homeodomain of UNCX proteins from different species.

## Notes

http://endoatlastest-env.eba-qtpfn7qz.us-east-1.elasticbeanstalk.com/

### Competing Interest Statement

The authors have declared no competing interest.

## REFERENCES AND NOTES

Achim, K., Pettit, J.B., Saraiva, L.R., Gavriouchkina, D., Larsson, T., Arendt, D., and Marioni, J.C. (2015). High-throughput spatial mapping of single-cell RNA-seq data to tissue of origin. Nat Biotechnol 33, 503–509. 10.1038/nbt.3209.

Bateson, W. (1894). Materials for the study of variation treated with especial regard to discontinuity in the origin of species (Macmillan and Company).

Benazeraf, B., and Pourquie, O. (2013). Formation and segmentation of the vertebrate body axis. Annu Rev Cell Dev Biol 29, 1–26. 10.1146/annurev-cellbio-101011-155703.

Berking, S. (2007). Generation of bilateral symmetry in Anthozoa: a model. J Theor Biol 246, 477–490. 10.1016/j.jtbi.2007.01.008.

Burtscher, I., and Lickert, H. (2009). Foxa2 regulates polarity and epithelialization in the endoderm germ layer of the mouse embryo. Development 136, 1029–1038. 10.1242/dev.028415.

Chen, J.A., Voigt, J., Gilchrist, M., Papalopulu, N., and Amaya, E. (2005). Identification of novel genes affecting mesoderm formation and morphogenesis through an enhanced large scale functional screen in Xenopus. Mech Dev 122, 307–331. 10.1016/j.mod.2004.11.008.

Cho, S.J., and Park, S.C. (2008). Paired-like subclass homeobox genes from the clitellate annelid Perionyx excavatus. Biochem Genet 46, 737–743. 10.1007/s10528-008-9189-z.

Chourrout, D., Delsuc, F., Chourrout, P., Edvardsen, R.B., Rentzsch, F., Renfer, E., Jensen, M.F., Zhu, B., de Jong, P., Steele, R.E., and Technau, U. (2006). Minimal ProtoHox cluster inferred from bilaterian and cnidarian Hox complements. Nature 442, 684–687. 10.1038/nature04863.

Christ, B., Schmidt, C., Huang, R., Wilting, J., and Brand-Saberi, B. (1998). Segmentation of the vertebrate body. Anat Embryol (Berl) 197, 1–8. 10.1007/s004290050116.

De Graeve, F., Jagla, T., Daponte, J.P., Rickert, C., Dastugue, B., Urban, J., and Jagla, K. (2004). The ladybird homeobox genes are essential for the specification of a subpopulation of neural cells. Dev Biol 270, 122–134. 10.1016/j.ydbio.2004.02.014.

Deng, M., Wang, Y., Zhang, L., Yang, Y., Huang, S., Wang, J., Ge, H., Ishibashi, T., and Yan, Y. (2019). Single cell transcriptomic landscapes of pattern formation, proliferation and growth in Drosophila wing imaginal discs. Development 146. 10.1242/dev.179754.

Diaz-Cuadros, M., Pourquie, O., and El-Sherif, E. (2021). Patterning with clocks and genetic cascades: Segmentation and regionalization of vertebrate versus insect body plans. PLoS Genet 17, e1009812. 10.1371/journal.pgen.1009812.

Donovan, K.M., Leidinger, M.R., McQuillen, L.P., Goeken, J.A., Hogan, C.M., Harwani, S.C., Flaherty, H.A., and Meyerholz, D.K. (2018). Allograft Inflammatory Factor 1 as an Immunohistochemical Marker for Macrophages in Multiple Tissues and Laboratory Animal Species. Comp Med 68, 341–348. 10.30802/AALAS-CM-18-000017.

Dray, N., Tessmar-Raible, K., Le Gouar, M., Vibert, L., Christodoulou, F., Schipany, K., Guillou, A., Zantke, J., Snyman, H., Behague, J., et al. (2010). Hedgehog signaling regulates segment formation in the annelid Platynereis. Science 329, 339–342. 10.1126/science.1188913.

Dries, R., Chen, J., Del Rossi, N., Khan, M.M., Sistig, A., and Yuan, G.C. (2021). Advances in spatial transcriptomic data analysis. Genome Res 31, 1706–1718. 10.1101/gr.275224.121.

Fritzenwanker, J.H., Saina, M., and Technau, U. (2004). Analysis of forkhead and snail expression reveals epithelial-mesenchymal transitions during embryonic and larval development of Nematostella vectensis. Dev Biol 275, 389–402. 10.1016/j.ydbio.2004.08.014.

Genikhovich, G., Fried, P., Prunster, M.M., Schinko, J.B., Gilles, A.F., Fredman, D., Meier, K., Iber, D., and Technau, U. (2015). Axis Patterning by BMPs: Cnidarian Network Reveals Evolutionary Constraints. Cell Rep 10, 1646–1654. 10.1016/j.celrep.2015.02.035.

He, S., Del Viso, F., Chen, C.Y., Ikmi, A., Kroesen, A.E., and Gibson, M.C. (2018). An axial Hox code controls tissue segmentation and body patterning in Nematostella vectensis. Science 361, 1377–1380. 10.1126/science.aar8384.

Hudry, B., Thomas-Chollier, M., Volovik, Y., Duffraisse, M., Dard, A., Frank, D., Technau, U., and Merabet, S. (2014). Molecular insights into the origin of the Hox-TALE patterning system. Elife 3, e01939. 10.7554/eLife.01939.

Hyman, L.H. (1940). The invertebrates: Protozoa through Ctenophora vol. 1 (The McGraw-Hill Companies).

Ikmi, A., and Gibson, M.C. (2010). Identification and in vivo characterization of NvFP-7R, a developmentally regulated red fluorescent protein of Nematostella vectensis. PLoS One 5, e11807. 10.1371/journal.pone.0011807.

Ikmi, A., Steenbergen, P.J., Anzo, M., McMullen, M.R., Stokkermans, A., Ellington, L.R., and Gibson, M.C. (2020). Feeding-dependent tentacle development in the sea anemone Nematostella vectensis. Nat Commun 11, 4399. 10.1038/s41467-020-18133-0.

Jagla, K., Dolle, P., Mattei, M.G., Jagla, T., Schuhbaur, B., Dretzen, G., Bellard, F., and Bellard, M. (1995). Mouse Lbx1 and human LBX1 define a novel mammalian homeobox gene family related to the Drosophila lady bird genes. Mech Dev 53, 345–356. 10.1016/0925-4773(95)00450-5.

Jagla, K., Jagla, T., Heitzler, P., Dretzen, G., Bellard, F., and Bellard, M. (1997). ladybird, a tandem of homeobox genes that maintain late wingless expression in terminal and dorsal epidermis of the Drosophila embryo. Development 124, 91–100. 10.1242/dev.124.1.91.

Juarez-Morales, J.L., Weierud, F., England, S.J., Demby, C., Santos, N., Grieb, G., Mazan, S., and Lewis, K.E. (2021). Evolution of lbx spinal cord expression and function. Evol Dev 23, 404–422. 10.1111/ede.12387.

Kamm, K., Schierwater, B., Jakob, W., Dellaporta, S.L., and Miller, D.J. (2006). Axial patterning and diversification in the cnidaria predate the Hox system. Curr Biol 16, 920–926. 10.1016/j.cub.2006.03.036.

Karaiskos, N., Wahle, P., Alles, J., Boltengagen, A., Ayoub, S., Kipar, C., Kocks, C., Rajewsky, N., and Zinzen, R.P. (2017). The Drosophila embryo at single-cell transcriptome resolution. Science 358, 194–199. 10.1126/science.aan3235.

Knabl, P., Schauer, A., Pomreinke, A.P., Zimmermann, B., Rogers, K.W., Müller, P., and Genikhovich, G. (2022). Analysis of SMAD1/5 target genes in a sea anemone reveals ZSWIM4-6 as a novel BMP signaling modulator. bioRxiv, 2022.2006.2003.494682. 10.1101/2022.06.03.494682.

Koch, T.L., and Grimmelikhuijzen, C.J.P. (2019). Global Neuropeptide Annotations From the Genomes and Transcriptomes of Cubozoa, Scyphozoa, Staurozoa (Cnidaria: Medusozoa), and Octocorallia (Cnidaria: Anthozoa). Front Endocrinol (Lausanne) 10, 831. 10.3389/fendo.2019.00831.

Kraus, Y., Aman, A., Technau, U., and Genikhovich, G. (2016). Pre-bilaterian origin of the blastoporal axial organizer. Nat Commun 7, 11694. 10.1038/ncomms11694.

Kusakabe, R., Higuchi, S., Tanaka, M., Kadota, M., Nishimura, O., and Kuratani, S. (2020). Novel developmental bases for the evolution of hypobranchial muscles in vertebrates. BMC Biol 18, 120. 10.1186/s12915-020-00851-y.

Lacin, H., Williamson, W.R., Card, G.M., Skeath, J.B., and Truman, J.W. (2020). Unc-4 acts to promote neuronal identity and development of the take-off circuit in the Drosophila CNS. Elife 9. 10.7554/eLife.55007.

Leclere, L., and Rentzsch, F. (2014). RGM regulates BMP-mediated secondary axis formation in the sea anemone Nematostella vectensis. Cell Rep 9, 1921–1930. 10.1016/j.celrep.2014.11.009.

Leitges, M., Neidhardt, L., Haenig, B., Herrmann, B.G., and Kispert, A. (2000). The paired homeobox gene Uncx4.1 specifies pedicles, transverse processes and proximal ribs of the vertebral column. Development 127, 2259–2267. 10.1242/dev.127.11.2259.

Lohoff, T., Ghazanfar, S., Missarova, A., Koulena, N., Pierson, N., Griffiths, J.A., Bardot, E.S., Eng, C.L., Tyser, R.C.V.g, Argelaguet, R., et al. (2022). Integration of spatial and single-cell transcriptomic data elucidates mouse organogenesis. Nat Biotechnol 40, 74–85. 10.1038/s41587-021-01006-2.

Mankoo, B.S., Skuntz, S., Harrigan, I., Grigorieva, E., Candia, A., Wright, C.V., Arnheiter, H., and Pachnis, V. (2003). The concerted action of Meox homeobox genes is required upstream of genetic pathways essential for the formation, patterning and differentiation of somites. Development 130, 4655–4664. 10.1242/dev.00687.

Mansouri, A., Voss, A.K., Thomas, T., Yokota, Y., and Gruss, P. (2000). Uncx4.1 is required for the formation of the pedicles and proximal ribs and acts upstream of Pax9. Development 127, 2251–2258. 10.1242/dev.127.11.2251.

Martindale, M.Q., Pang, K., and Finnerty, J.R. (2004). Investigating the origins of triploblasty: ‘mesodermal’ gene expression in a diploblastic animal, the sea anemone Nematostella vectensis (phylum, Cnidaria; class, Anthozoa). Development 131, 2463–2474. 10.1242/dev.01119.

Martinez Barbera, J.P., Clements, M., Thomas, P., Rodriguez, T., Meloy, D., Kioussis, D., and Beddington, R.S. (2000). The homeobox gene Hex is required in definitive endodermal tissues for normal forebrain, liver and thyroid formation. Development 127, 2433–2445. 10.1242/dev.127.11.2433.

Marx, V. (2021). Method of the Year: spatially resolved transcriptomics. Nat Methods 18, 9–14. 10.1038/s41592-020-01033-y.

Miller, D.M., Shen, M.M., Shamu, C.E., Burglin, T.R., Ruvkun, G., Dubois, M.L., Ghee, M., and Wilson, L. (1992). C. elegans unc-4 gene encodes a homeodomain protein that determines the pattern of synaptic input to specific motor neurons. Nature 355, 841–845. 10.1038/355841a0.

Miyasaka, K.Y., Kida, Y.S., Sato, T., Minami, M., and Ogura, T. (2007). Csrp1 regulates dynamic cell movements of the mesendoderm and cardiac mesoderm through interactions with Dishevelled and Diversin. Proc Natl Acad Sci U S A 104, 11274–11279. 10.1073/pnas.0702000104.

Moriel, N., Senel, E., Friedman, N., Rajewsky, N., Karaiskos, N., and Nitzan, M. (2021). NovoSpaRc: flexible spatial reconstruction of single-cell gene expression with optimal transport. Nat Protoc 16, 4177–4200. 10.1038/s41596-021-00573-7.

Neidhardt, L.M., Kispert, A., and Herrmann, B.G. (1997). A mouse gene of the paired-related homeobox class expressed in the caudal somite compartment and in the developing vertebral column, kidney and nervous system. Dev Genes Evol 207, 330–339. 10.1007/s004270050120.

Nguyen, H.T., Bodmer, R., Abmayr, S.M., McDermott, J.C., and Spoerel, N.A. (1994). D-mef2: a Drosophila mesoderm-specific MADS box-containing gene with a biphasic expression profile during embryogenesis. Proc Natl Acad Sci U S A 91, 7520–7524. 10.1073/pnas.91.16.7520.

Nitzan, M., Karaiskos, N., Friedman, N., and Rajewsky, N. (2019). Gene expression cartography. Nature 576, 132–137. 10.1038/s41586-019-1773-3.

Noden, D.M., Marcucio, R., Borycki, A.G., and Emerson, C.P., Jr. (1999). Differentiation of avian craniofacial muscles: I. Patterns of early regulatory gene expression and myosin heavy chain synthesis. Dev Dyn 216, 96–112. 10.1002/(SICI)1097-0177(199910)216:2<96::AID-DVDY2>3.0.CO;2-6.

Ochi, H., and Westerfield, M. (2009). Lbx2 regulates formation of myofibrils. BMC Dev Biol 9, 13. 10.1186/1471-213X-9-13.

Onai, T., Irie, N., and Kuratani, S. (2014). The evolutionary origin of the vertebrate body plan: the problem of head segmentation. Annu Rev Genomics Hum Genet 15, 443–459. 10.1146/annurev-genom-091212-153404.

Pax, F. (1913). Die Actinien (G. Plaetzsche Buchdruckerei Lippert & Company).

Pedersen, J.K., Nelson, S.B., Jorgensen, M.C., Henseleit, K.D., Fujitani, Y., Wright, C.V., Sander, M., Serup, P., and Beta Cell Biology, C. (2005). Endodermal expression of Nkx6 genes depends differentially on Pdx1. Dev Biol 288, 487–501. 10.1016/j.ydbio.2005.10.001.

Pourquie, O. (2000). Segmentation of the paraxial mesoderm and vertebrate somitogenesis. Curr Top Dev Biol 47, 81–105. 10.1016/s0070-2153(08)60722-x.

Rentzsch, F., Anton, R., Saina, M., Hammerschmidt, M., Holstein, T.W., and Technau, U. (2006). Asymmetric expression of the BMP antagonists chordin and gremlin in the sea anemone Nematostella vectensis: implications for the evolution of axial patterning. Dev Biol 296, 375–387. 10.1016/j.ydbio.2006.06.003.

Rodriques, S.G., Stickels, R.R., Goeva, A., Martin, C.A., Murray, E., Vanderburg, C.R., Welch, J., Chen, L.M., Chen, F., and Macosko, E.Z. (2019). Slide-seq: A scalable technology for measuring genome-wide expression at high spatial resolution. Science 363, 1463–1467. 10.1126/science.aaw1219.

Ryan, J.F., Mazza, M.E., Pang, K., Matus, D.Q., Baxevanis, A.D., Martindale, M.Q., and Finnerty, J.R. (2007). Pre-bilaterian origins of the Hox cluster and the Hox code: evidence from the sea anemone, Nematostella vectensis. PLoS One 2, e153. 10.1371/journal.pone.0000153.

Satija, R., Farrell, J.A., Gennert, D., Schier, A.F., and Regev, A. (2015). Spatial reconstruction of single-cell gene expression data. Nat Biotechnol 33, 495–502. 10.1038/nbt.3192.

Saudemont, A., Dray, N., Hudry, B., Le Gouar, M., Vervoort, M., and Balavoine, G. (2008). Complementary striped expression patterns of NK homeobox genes during segment formation in the annelid Platynereis. Dev Biol 317, 430–443. 10.1016/j.ydbio.2008.02.013.

Schragle, J., Huang, R., Christ, B., and Prols, F. (2004). Control of the temporal and spatial Uncx4.1 expression in the paraxial mesoderm of avian embryos. Anat Embryol (Berl) 208, 323–332. 10.1007/s00429-004-0404-3.

Scott, M.P., and Carroll, S.B. (1987). The segmentation and homeotic gene network in early Drosophila development. Cell 51, 689–698. 10.1016/0092-8674(87)90092-4.

Seaver, E.C. (2003). Segmentation: mono-or polyphyletic? Int J Dev Biol 47, 583–595.

Sebe-Pedros, A., Saudemont, B., Chomsky, E., Plessier, F., Mailhe, M.P., Renno, J., Loe-Mie, Y., Lifshitz, A., Mukamel, Z., Schmutz, S., et al. (2018). Cnidarian Cell Type Diversity and Regulation Revealed by Whole-Organism Single-Cell RNA-Seq. Cell 173, 1520–1534 e1520. 10.1016/j.cell.2018.05.019.

Steinmetz, P.R.H. (2019). A non-bilaterian perspective on the development and evolution of animal digestive systems. Cell Tissue Res 377, 321–339. 10.1007/s00441-019-03075-x.

Steinmetz, P.R.H., Aman, A., Kraus, J.E.M., and Technau, U. (2017). Gut-like ectodermal tissue in a sea anemone challenges germ layer homology. Nat Ecol Evol 1, 1535–1542. 10.1038/s41559-017-0285-5.

Stickels, R.R., Murray, E., Kumar, P., Li, J., Marshall, J.L., Di Bella, D.J., Arlotta, P., Macosko, E.Z., and Chen, F. (2021). Highly sensitive spatial transcriptomics at near-cellular resolution with Slide-seqV2. Nat Biotechnol 39, 313–319. 10.1038/s41587-020-0739-1.

Takahashi, T. (2020). Comparative Aspects of Structure and Function of Cnidarian Neuropeptides. Front Endocrinol (Lausanne) 11, 339. 10.3389/fendo.2020.00339.

Tautz, D. (2004). Segmentation. Dev Cell 7, 301–312. 10.1016/j.devcel.2004.08.008.

Technau, U. (2020). Gastrulation and germ layer formation in the sea anemone Nematostella vectensis and other cnidarians. Mech Dev 163, 103628. 10.1016/j.mod.2020.103628.

Treffkorn, S., Kahnke, L., Hering, L., and Mayer, G. (2018). Expression of NK cluster genes in the onychophoran Euperipatoides rowelli: implications for the evolution of NK family genes in nephrozoans. Evodevo 9, 17. 10.1186/s13227-018-0105-2.

van den Brink, S.C., Alemany, A., van Batenburg, V., Moris, N., Blotenburg, M., Vivie, J., Baillie-Johnson, P., Nichols, J., Sonnen, K.F., Martinez Arias, A., and van Oudenaarden, A. (2020). Single-cell and spatial transcriptomics reveal somitogenesis in gastruloids. Nature 582, 405–409. 10.1038/s41586-020-2024-3.

Walthall, W.W. (1995). Repeating patterns of motoneurons in nematodes: the origin of segmentation? EXS 72, 61–75. 10.1007/978-3-0348-9219-3_4.

Watanabe, H., Kuhn, A., Fushiki, M., Agata, K., Ozbek, S., Fujisawa, T., and Holstein, T.W. (2014). Sequential actions of beta-catenin and Bmp pattern the oral nerve net in Nematostella vectensis. Nat Commun 5, 5536. 10.1038/ncomms6536.

Wijesena, N., Simmons, D.K., and Martindale, M.Q. (2017). Antagonistic BMP-cWNT signaling in the cnidarian Nematostella vectensis reveals insight into the evolution of mesoderm. Proc Natl Acad Sci U S A 114, e5608–E5615. 10.1073/pnas.1701607114.

Wilm, B., James, R.G., Schultheiss, T.M., and Hogan, B.L. (2004). The forkhead genes, Foxc1 and Foxc2, regulate paraxial versus intermediate mesoderm cell fate. Dev Biol 271, 176–189. 10.1016/j.ydbio.2004.03.034.

Wotton, K.R., Weierud, F.K., Dietrich, S., and Lewis, K.E. (2008). Comparative genomics of Lbx loci reveals conservation of identical Lbx ohnologs in bony vertebrates. BMC Evol Biol 8, 171. 10.1186/1471-2148-8-171.

