## Supplemental Information for "Spatial transcriptomics reveals a conserved segment polarity program that governs muscle patterning in *Nematostella vectensis*"

### CONTACT FOR REAGENT AND RESOURCE SHARING

### EXPERIMENTAL MODEL AND SUBJECT DETAILS

Animal Husbandry, spawning induction and microinjection

*Nematostella* colonies were cultured at 22°C in 12 parts per thousand (ppt) artificial seawater (ASW; Sea Salt from Instant Ocean) until reaching sexual maturity. Spawning induction, de-jellying and microinjection were carried out as following published protocols (Genikhovich and Technau, 2009c; Stefanik et al., 2013). Embryos were cultured at 22°C following fertilization.

### METHOD DETAILS

Immunohistochemistry and Fluorescent *in situ* hybridization (FISH)

Animals were fixed and stained following previously published protocols (Genikhovich and Technau, 2009a; Wolenski et al., 2013). In brief, planula larvae and polyps were anesthetized by dissolving 7% $MgCl_2$ in 12ppt ASW. Animals used for FISH were pre-fixed at room temperature in 4% paraformaldehyde, 2.5% glutaraldehyde in 12ppt ASW for 90s. Animals were then fixed in 4% Paraformaldehyde in 12ppt ASW for 1 hour at room temperature. The following antibodies and dyes were used for immunostaining: SiR-actin (Cytoskeleton, 1:1000), SiR-DNA (Cytoskeleton, 1:1000), SYBR-Green (ThermoFisher, 1:10000), anti-NvVasa1 (citation, 1:1000). FISH was carried out following a published protocol (Genikhovich and Technau, 2009b; Wolenski *et al.*, 2013). Animals were hybridized with 1µg of DIG-labeled RNA probes at 60°C for 48 hours and incubated in pre-absorbed anti-DIG-POD fab fragment (Roche, 1:1000)/5% sheep serum (Sigma)/1% blocking reagent (Roche)/TNT buffer at 4°C overnight. Fluorescent labeling was performed using the TSA Cyanine 3 (Cy3) detection kit (Akoya) for 20 min at RT. Animals were then mounted in ScaleA2 buffer (2M Urea, 75% Glycerol) with desired angle and imaged.

Imaging and image quantification

Animals were imaged using either Leica SP5 confocal microscope with LAS AF software or Zeiss 700 upright confocal microscope with ZEN software. All FISH samples were imaged under the same setting. Acquired images were processed using Fiji. The angle tool was utilized for segment angle measurements and "Plot profile" function was utilized to obtain intensity plot for FISH images. Raw measurements can be downloaded from the Stowers Institute for Medical Research Original Data Repository.

CRISPR/Cas9-mediated mutagenesis

CRISPR/Cas9 genome editing in *Nematostella* embryos was performed as previously described (Ikmi et al., 2014). GuideRNAs were designed using the CRISPRscan online web interface (http://www.crisprscan.org/?page=sequence)(Moreno-Mateos et al., 2015) with default settings (PAM= NGG, in vitro T7 promoter). To generate double stranded DNA templates for gRNAs, the gene specific forward primers were first annealed to the universal reverse primer and underwent Klenow extension (37°C 30 mins, 75°C 10mins, Klenow fragment, NEB, 1:20). In vitro transcription was carried out using the AmpliScribe™ T7-Flash™ Transcription Kit (Epicenter) at 37°C for 4 hours. gRNAs were purified using the Direct-zol™ RNA MiniPrep Kit (Zymo Research) following manufacture instructions and quantified with a NanoDrop spectrophotometer (ThermoFisher Scientific). Prior to injection, SpCas9 protein with NLS sequence (PNA Bio) was mixed with equal concentration of gRNA and diluted to 500ng/µL each using DEPC water. Injected oocytes were fertilized with wild-type sperm and raised at 22°C. Aboral amputations were made to obtain tissue fragments from individual animal for genotyping. Tissue was lysed in QuickExtract™ DNA Extraction Solution (Lucigen) at 50°C for 30 mins and 98°C for 2 mins before PCR. All primers used for gRNA synthesis and genotyping can be found in the Materials section.

shRNA-mediated gene knockdown

Short hairpin RNA (shRNA) mimics precursor microRNA (pre-miRNA) during endogenous miRNA processing and has been widely applied in mammalian tissue culture systems. Here, we optimized the conventional plasmid-based shRNA approach to better suit the biological feature of *Nematostella* embryos.

For shRNA design, sequences of genes of interest were submitted to siRNA Wizard web interface (InvivoGen; http://www.invivogen.com/sirnawizard/design.php) with motif size of 19nt. Candidate sequences with high complexity as well as optimal GC content of around 50% were selected and BLASTed against *Nematostella* transcriptome to ensure their specificity. The full-length sequence of shRNA, including forward and reverse stem sequence, separated by a 9nt linker (5'-TTCAAGAGA-3') was then submitted to the Mfold Web Server (RNA Institute, State University of New York at Albany; http://unafold.rna.albany.edu/?q=mfold/RNA-Folding-Form) to ensure the formation of stable stem-loop structures in vitro. Partially overlapping forward and reverse primers were ordered from Integrated DNA Technologies, and PCR amplified to get the 66nt DNA template, containing T7 promoter and the full-length shRNA sequence. Alternatively, full-length forward and reverse primers can be annealed to directly synthesize the template. Two extra thymidines (T) were added on the 3-prime end of the template to mimic endogenous pre-miRNA structure. In vitro transcription was performed using AmpliScribe™ T7-Flash™ Transcription Kit (Epicenter). shRNAs were purified using Direct-zol™ RNA MiniPrep Kit (Zymo Research) and quantified using NanoDrop spectrophotometer (ThermoFisher Scientific).

In vitro synthesized shRNAs were delivered into unfertilized *Nematostella* oocytes via microinjection. shRNA working concentration is ranged from 200ng/µL to 500ng/µL, depending on the expression level and developmental stage that the gene is expressed. In general, the higher the expression level, or the later the developmental stage, the more shRNA is needed. shRNAs were diluted to desired concentrations and co-injected with Dextran Red (0.2 µg/µl, Molecular Probes) as a long-term tracer to distinguish un-injected animals. Oocytes were then fertilized with fresh wild-type sperm and reared at 22°C after injection.

Endomesoderm isolation and single cell RNA-seq

To obtain endomesoderm isolates, wildtype embryos were raised at 22°C with daily water changes until 72hpf in a glass dish. Around 200 planula larvae were manually selected based on their gross morphology (no excess tissue, no axis duplication and show clear signs of endomesodermal segmentation) and transferred to a gelatin-coated 35x10 mm Falcon petri dish containing 1% Sodium Thioglycolate, 50mM NaCl, 5mM EDTA in 12ppt $Ca^{2+}$/$Mg^{2+}$ free ASW (CMFSW). The planula larvae were incubated for 7 mins at room temperature with gentle shaking and then transferred into a new gelatin-coated petri dish containing fresh 12ppt CMFSW. With gentle pipetting, the ectodermal tissue will start to peel off, leaving the intact endomesoderm and partial pharyngeal tissue encapsulated by the mesoglea. The endomesoderm isolates were then carefully transferred using a gelatin-coated 200µL pipette tip into a pre-chilled 1.5mL Eppendorf tube. 1mL ice cold StemPro® Accutase® Cell Dissociation Reagent (ThermoFisher) was then added to the tube and incubated for 10 mins at room temperature. Gentle pipetting was applied to increase the rate of dissociation. Next, the lysate was passed through a Flowmi® Cell Strainer (Sigma) with porosity of 40µm to remove mesoglea and un-dissociated cell clumps. This lysate was then centrifuged at 300 rpm for 5 mins at 4°C. Cell palette was resuspended in 200µL ice cold 1x PBS with 1% BSA and immediately proceeded to viability check.

Library was constructed following the protocol from Chromium Single Cell 3' Kit (version 3.1, 10X Genomics). Sequencing was performed using Illumina NovaSeq system with 75bp paired reads.

single cell mRNA-seq analysis

10X scRNA-seq library was sequenced via Illumina Novaseq 6000 instrument using the following paired end length: 28 bp Read1, 8 bp I7 Index and 98 bp Read2. Raw reads were processed using 10X Genomics pipeline Cellranger (version 4.0). Reads were demultiplexed into fastq file format using function *cellranger mkfastq*. Genome index was built by cellranger mkref using nematostella genome Nvec200. Fastq files were aligned by STAR aligner and cell UMI counts table were generated using cellranger *count* function with default parameters.

scRNAseq signals were analyzed with Seurat, only cells with more than 200 detected genes were retained for downstream analysis. Cell clustering was performed using the top 1500 most variable features and visualized using UMAP. Cell clusters were annotated based on the top 10 marker genes per cluster as well as additional published marker genes. Table S1 summarized the information of all the genes used in this manuscript, their corresponding annotations, and gene IDs in two publicly available *Nematostella* transcriptomes (NV2 and UVienna).

Bulk RNA-seq and identification of landmark genes

*Anthox1a*, *Anthox8* and *Anthox6a* F3 homozygous mutants were generated as previously described (He et al., 2018). Sexually mature F2 homozygous animals were kept at 18°C in a dark incubator for two weeks and then light shocked to induce spawning. Homozygous F3 embryos were raised at 22°C and harvested at 48, 60, 72hpf, respectively. 200 larvae were collected for each mutant strain per replica per time point.

For the *Pbx* knockdown experiment, two independent shRNAs targeting *Pbx* were injected into unfertilized oocytes and fertilized with wildtype sperm. Injected animals were raised at 22°C and harvested 72hpf. 200 larvae were collected per replica.

Total RNA was extracted using Direct-zol™ RNA MiniPrep Kit (Zymo Research) following manufacture instructions and quantified using QubitTM RNA HS Assay Kit (ThermoFisher).

After removing adapter sequences with cutdapt (v 2.10), RNA-seq reads were trimmed to 50bp before aligning to the NV2 genome with STAR (v 2.7.1a). The number of uniquely mapped reads at annotated genes were then quantified using featureCounts (v 1.6.3). RUVr (k=1) was performed to minimize batch effect and the resulting normalized signals were used for PCA and differential expression analysis. Differentially expressed genes (DEGs, absolute foldchange >=2 and adjusted pvalue <= 0.01) under each Hox mutant condition at each time point were identified using DESeq2. With a particular focus on transcription factors and signaling pathway components, 151 candidate gene were selected from the DEG list for in situ validation. Out of these 151 genes, 51 displayed restricted yet stereotypic expression patterns in the endomesoderm of 72hpf planula larvae and were used as landmark genes.

<u>Construction of Endo-atlas</u>

To construct Endo-atlas, we performed spatial expression reconstruction using novoSpaRc (Nitzan et al., 2019). In short, novoSpaRc employs the assumption that cells that are physically close tend to share similar transcription profiles and infers the probabilities of spatial position for individual cells using the spatial expression pattern of reference marker genes. Within the framework of novoSpaRc, we first selected 34 marker genes whose spatial expression patterns were identified with *in situ* hybridization. We then constructed a virtual 3D model of *Nematostella* embryo endomesoderm using blender® and converted the expression profile of our marker genes to binarized values at 8993 locations along the surface of this 3D model. Next, we selected 2623 cells with expression of at least 5 of our marker genes from scRNAseq data and inferred the spatial expression profile of the other genes using novoSpaRc marker guided reconstruction (with alpha = 0.5). The predicted expression patterns were then visualized in 3D using plotly.

Phylogenetic analysis

For phylogenetic analysis of *Lbx*, the following LBX protein sequences were downloaded from NCBI: *Homo sapiens* LBX1 (NP_006553.2), LBX2 (NP_001269359.1); *Mus musculus* LBX1 (NP_034821.2), LBX2 (NP_034822.1); *Xenopus laevis* LBX1 (NP_001089192.1); *Rattus norvegicus* LBX1 (NP_001040573.1), LBX2 (NP_001102714.1), *Gallus gallus* LBX1 (NP_001171614.1), LBX2 (NP_001038133.1); *Danio rerio* LBX1a (NP_001020703.1), LBX1b (NP_001156784.1), LBX2 (NP_001007135.1); *Oikopleura dioica* LBX (AAW23077.1); *Branchiostoma floridae* LBX (XP_035672500.1); *Drosophila melanogaster* LBE (ACN74238.1), LBL (ACN74522.1); *Tribolium castaneum* LBX (XM_969913.3); *Apis cerana* LBX (A0A2A3EQ26); *Stylophora pistillata* LBX (PFX30300.1); *Amphimedon queenslandica* HOXB4-like (NP_001292180.1). *Acropora digitifera* LBX (XP_015770678.1) sequence was downloaded from OIST Marine Genomics Unit (https://marinegenomics.oist.jp/adig/viewer/info?project_id=87, Shinzato et al., 2021).

For phylogenetic analysis of Uncx, the following UNCX protein sequences were downloaded from NCBI: *Homo sapiens* UNCX (NP_001073930.1); *Mus musculus* UNCX (NP_038730.1), *Rattus norvegicus* UNCX (NP_058875.2), *Gallus gallus* UNCX (XP_015149815.1); *Danio rerio* UNCX (XP_005164261.1), UNCX4.1 (NP_001018616.2); *Xenopus tropicalis* UNCX (NP_001305673.1); *Drosophila melanogaster* UNC-4 (NP_573242.2); *Tribolium castaneum* UNC-4 (XP_969276.2); *Argiope bruennichi* UNC-4 (KAF8793013.1); *Caenorhabditis elegans* UNC-4 (NP_496138.1); *Clytia hemisphaerica* UNC (partial protein, ALJ33549.1); *Amphimedon queenslandica* RAX-like (XP_003383252.1). *Acropora digitifera* UNCX (aug_v2a.06347.t1) sequence was downloaded from OIST Marine Genomics Unit (https://marinegenomics.oist.jp/adig/viewer/info?project_id=87, Shinzato *et al.*, 2021). *Montipora capitata* UNCX (augustus.g46304.t1) was downloaded from http://cyanophora.rutgers.edu/montipora (Shumaker et al., 2019).

The web interface of EMBL-EBI Clustal Omega (https://www.ebi.ac.uk/Tools/msa/clustalo/, McWilliam et al., 2013; Sievers et al., 2011) was used to align the sequences and maximum likelihood tree was constructed using IQ-TREE (http://www.iqtree.org/, Minh et al., 2020; Nguyen et al., 2015) using the following parameters, bootstrap replicates = 1000, perturbation strength = 0.5, unsuccessful iterations to stop = 100. 'ggtree' package was used to plot the inferred phylogeny in R (version 4.1.3).

### DATA AND SOFTWARE AVAILABILITY

Original data underlying this manuscript can be downloaded from Stowers Institute for Medical Research Original Data Repository: http://www.stowers.org/research/publications/libpb-1623. All NGS sequencing results can be downloaded from the GEO repository at: https://www.ncbi.nlm.nih.gov/geo/query/acc.cgi?acc=GSE173786. The *Nematostella* Endo-atlas can be accessed at: http://endoatlastest-env.eba-qtpfn7qz.us-east-1.elasticbeanstalk.com/. The NV2 genome and transcriptome used in this study are hosted on SIMRbase: https://genomes.stowers.org/.

### Table S1. Annotations, gene IDs and references of all the genes mentioned in this study.

| Gene Name | Protein annotation | Gene ID (NV2) | Gene ID (UVienna) | Reference |
|---|---|---|---|---|
| *Actb* | Actin, cytoplasmic 1 | NV2g013196000.1 | NVE14201 | |
| *Adamts7* | A disintegrin and metalloproteinase with thrombospondin motifs 7 | NV2g008406000.1 | NVE7526, NVE23888 | |
| *Aebp1* | Adipocyte enhancer-binding protein 1 | NV2g017174000.1 | NVE25694 | |
| *Agk* | Arginine kinase | NV2g017940000.1 | NVE16356 | |
| *Alk* | ALK tyrosine kinase receptor | NV2g006785000.1 | NVE25858 | |
| *Alx4* | Homeobox protein aristaless-like 4 | NV2g017442000.1 | NVE1403 | |
| *Anthox1* | Homeobox protein Anthox1 | NV2g017961000.1 | NVE16373 | (He *et al.*, 2018; Ryan et al., 2007) |
| *Anthox1a* | Homeobox protein Anthox1a | NV2g012279000.1 | NVE12998 | (He *et al.*, 2018; Ryan *et al.*, 2007) |
| *Anthox6a* | Homeobox protein Anthox6a | NV2g011327000.1 | NVE11345 | (He *et al.*, 2018; Ryan *et al.*, 2007) |
| *Anthox7* | Homeobox protein Anthox7 | NV2g012004000.1 | NVE21278 | (He *et al.*, 2018; Ryan *et al.*, 2007) |
| *Anthox8a* | Homeobox protein Anthox8a | NV2g012007000.1 | NVE21156 | (He *et al.*, 2018; Ryan *et al.*, 2007) |
| *Anthox8b* | Homeobox protein Anthox8b | NV2g012008000.1 | NVE21279 | (He *et al.*, 2018) |
| *Aplp* | Apolipophorin | NV2g000914000.1 | NVE25290, NVE3548 | |
| *Arp6* | Neurogenic differentiation factor 1 | NV2g006611000.1 | NVE5998 | (Watanabe et al., 2014) |
| *Art5* | Erythroblast NAD(P)(+)--arginine ADP-ribosyltransferase | NV2g012927000.1 | NVE12648, NVE12647 | |
| *Ascl5* | Achaete-scute homolog 5 | NV2g009665000.1 | NVE15559 | |
| *Asmt* | Acetylserotonin O-methyltransferase | NV2g016644000.1 | NVE2902 | |
| *Atoh1* | Protein atonal homolog 1 | NV2g010861000.1 | NVE18116 | |
| *B-H1* | Homeobox protein B-H1 | NV2g025278000.1 | NVE18480 | |
| *Bra(TbxT)* | Brachyury/T-box transcription factor T homolog 1 | NV2g010624000.1 | NVE3568 | (Scholz and Technau, 2003) |
| *Btbd16* | BTB/POZ domain-containing protein 16 | NV2g017664000.1 | NVE20145 | |
| *Cam* | Calmodulin | NV2g014600000.1 | NVE8332 | |
| *Casr* | Casr: Extracellular calcium-sensing receptor | NV2g025295000.1 NV2g025296000.1 | NVE18492 | |
| *Cdx* | HoxB3-like | NV2g030001000.1 | | |
| *Col27a1b* | Collagen alpha-1(XXVII) chain B | NV2g012027000.1 | NVE21289 | |
| *Col4a1* | Collagen alpha-1(IV) chain | NV2g010959000.1 | NVE19791, NVE19792, NVE7461 | |
| *Col6a5* | Collagen alpha-5(VI) chain | NV2g013883000.1 | NVE21880 | |
| *Cpa2* | Carboxypeptidase A2 | NV2g001253000.1 | NVE13106 | |
| *Cthrc1* | Collagen triple helix repeat-containing protein 1 | NV2g005041000.1 | NVE6626 | |
| *Ctrb2* | Chymotrypsinogen B2 | NV2g013528000.1 | NVE23810 | |
| *Ctsh* | Similar to Cathepsin L | NV2g004249000.1 | NVE5223 | |
| *D5-like* | D(5)-like dopamine receptor | NV2g012459000.1 | NVE17593 | |
| *DBH-like 1* | DBH-like monooxygenase protein 1 | NV2g013142000.1 | NVE14232, NVE5920 | |
| *Dlx* | Homeobox protein Dlx | NV2g014640000.1 | NVE8363 | (Ryan *et al.*, 2007) |

| | | | | |
|---|---|---|---|---|
| *Dmbx1* | Diencephalon/mesencephalon homeobox protein 1 | NV2g003591000.1 | NVE1803 | |
| *Dmbx4* | Diencephalon/mesencephalon homeobox protein 1-B | NV2g003592000.1 | NVE1799 | |
| *Dmbx5* | Diencephalon/mesencephalon homeobox protein 1-B | NV2g003593000.1 | NVE1797 | |
| *Efemp1* | EGF-containing fibulin-like extracellular matrix protein 1 | NV2g013293000.1 | NVE12199 | |
| *Elav* | ELAV-like protein | NV2g000252000.1 | NVE8968 | (Nakanishi et al., 2012) |
| *Elel1* | Similar to L-rhamnose-binding lectin ELEL-1 | NV2g008609000.1 | NVE13796 | |
| *Emx* | Homeobox protein EMX1 | NV2g018403000.1 | NVE4528 | |
| *F5* | F5: Coagulation factor V | NV2g000672000.1 | NVE6185, NVE5356 | |
| *F8* | F8: Coagulation factor VIII | NV2g004588000.1 | NVE10856 | |
| *Fbxw7* | F-box/WD repeat-containing protein 7 | NV2g005429000.1 | NVE7672, NVE12414 | |
| *Fgfrl* | Fibroblast growth factor receptor 4 | NV2g019093000.1 | NVE19166 | |
| *Fkbp14* | Peptidyl-prolyl cis-trans isomerase FKBP14 | NV2g011564000.1 | NVE14917 | |
| *FoxA* | Forkhead box protein A2 | NV2g011441000.1 | NVE20630 | (Fritzenwanker et al., 2004) |
| *FoxQ1* | Forkhead box protein Q1 | NV2g004570000.1 | NVE10869 | |
| *Fth* | Similar to Soma ferritin | NV2g019682000.1 | NVE15943 | |
| *Gbx* | Gastrulation brain homeodomain | NV2g011380000.1 | NVE20684 | (He *et al.*, 2018) |
| *Gdf5* | Growth/differentiation factor 6 | NV2g006489000.1 | NVE17522 | (Leclere and Rentzsch, 2014) |
| *Ggt* | Glutathione hydrolase 1 proenzyme | NV2g001925000.1 | NVE20501 | (Babonis and Martindale, 2017) |
| *Gp2* | Pancreatic secretory granule membrane major glycoprotein GP2 | NV2g014524000.1 | NVE9308 | |
| *Gremlin1* | Gremlin-1 | NV2g000122000.1 | NVE16132 | (Rentzsch et al., 2006) |
| *Gsc* | Homeobox protein goosecoid | NV2g010922000.1 | NVE19762 | (Matus et al., 2006) |
| *Hand1* | Heart- and neural crest derivatives-expressed protein | NV2g002675000.1 | NVE24171 | (Wijesena et al., 2017) |
| *Hand2* | Heart-and neural crest derivatives-expressed protein | NV2g002672000.1 | NVE24170 | (Wijesena *et al.*, 2017) |
| *Hmcn1* | Hemicentin-1 | NV2g023465000.1 | NVE10021 | |
| *Hpcal1* | Hippocalcin-like protein 1 | NV2g003389000.1 | NVE8130 | |
| *LactB2* | Beta-lactamase-like protein 2 | NV2g011772000.1 | NVE18995 | |
| *Lbx* | Transcription factor LBX2 | NV2g017667000.1 | NVE20146 | |
| *MelC4* | Myosin-2 essential light chain | NV2g005271000.1 | NVE3912 | (Cole et al., 2020) |
| *Mmp17* | Matrix metalloproteinase-17 | NV2g020474000.1 | NVE22474 | |
| *Mmp19* | Matrix metalloproteinase-19 | NV2g008375000.1 | NVE7504 | |
| *MoxB* | Homeobox protein Mox1 | NV2g012940000.1 | NVE10742 | (Ryan *et al.*, 2007) |
| *MoxC* | Homeobox protein Mox1 | NV2g012942000.1 | NVE10743 | (Ryan *et al.*, 2007) |
| *MoxD* | Homeobox protein Mox1 | NV2g012943000.1 | NVE10744 | (Ryan *et al.*, 2007) |
| *Msx1* | Homeobox protein MSX-1 | NV2g012247000.1 | NVE12977 | |
| *Myc* | Myc proto-oncogene protein | NV2g018825000.1 | NVE20961 | |
| *MyHC-st* | Similar to Myosin heavy chain, striated muscle | NV2g008024000.1 | NVE14552 | |
| *Naldl1* | Aminopeptidase NAALADL1 | NV2g011844000.1 | NVE18936 | |

| | | | | |
|---|---|---|---|---|
| *Ncol3* | Minicollagen 3 | NV2g005200000.1 | NVE3845 | (Babonis and Martindale, 2017) |
| *Ncol4* | Minicollagen 4 | NV2g011801000.1 | NVE18974 | (Babonis and Martindale, 2017) |
| *Netrin* | Netrin-1 | NV2g007734000.1 | NVE5022 | (Matus *et al.*, 2006) |
| *NeuroD1* | Neurogenic differentiation factor 1 | NV2g006608000.1 | NVE6000 | |
| *Ngal* | Nematogalectin | NV2g005200000.1 | NVE3845 | (Babonis and Martindale, 2017) |
| *Nkx2.2a1* | Homeobox protein Nkx-2.2a | NV2g011134000.1 | NVE10557 | (Steinmetz et al., 2017) |
| *Nkx2.2a2* | Homeobox protein Nkx-2.2a | NV2g011262000.1 | NVE11289 | (Steinmetz *et al.*, 2017) |
| *Nkx2.8* | Homeobox protein Nkx-2.8 | NV2g011132000.1 | NVE10555 | |
| *Nkx3.2* | Homeobox protein Nkx-3.2 | NV2g018723000.1 | NVE24919 | (Steinmetz *et al.*, 2017) |
| *Noggin* | Noggin | NV2g007394000.1 | | (Saina et al., 2009) |
| *Notum1* | Palmitoleoyl-protein carboxylesterase NOTUM | NV2g008355000.1 | NVE7485 | |
| *Notum2* | Palmitoleoyl-protein carboxylesterase NOTUM | NV2g025566000.1 | NVE1463, NVE1464 | |
| *NV2.10455* | Unknown | NV2g010455000.1 | NVE20336 | |
| *NV2.1690* | Unknown | NV2g001690000.1 | NVE478 | |
| *NV2.16999* | unknown | NV2g016999000.1 | | |
| *NV2.16999* | Unknown | NV2g016999000.1 | | |
| *NV2.18749* | TNF receptor-associated factor 6 | NV2g018749000.1 | NVE20902 | |
| *NV2.19138* | Unknown | NV2g019138000.1 | NVE17250 | |
| *NV2.23364* | Ankyrin repeat domain-containing protein 49 | NV2g023364000.1 | NVE12127 | |
| *NV2.2940* | Unknown | NV2g002940000.1 | NVE5451 | |
| *NV2.4713* | Unknown | NV2g004713000.1 | NVE825 | |
| *NV2.5420* | Unknwon | NV2g005420000.1 | NVE7659 | |
| *NV2.5900* | Unknown | NV2g005900000.1 | NVE12038 | |
| *NV2.7862* | Unknown | NV2g007862000.1 | NVE2716 | |
| *NV2.7863* | Unknown | NV2g007863000.1 | NVE2717 | |
| *NV2.9093* | Unknown | NV2g009093000.1 | NVE6814, NVE1643 | |
| *NV2.9262* | Unknown | NV2g009262000.1 | | |
| *OtxA* | Homeobox protein OTX1 | NV2g012793000.1 | NVE7117 | (Mazza et al., 2007) |
| *OtxB* | Homeobox protein OTX1 B | NV2g012795000.1 | NVE7116 | (Mazza *et al.*, 2007) |
| *OtxC* | Homeobox protein OTX1 | NV2g012799000.1 | NVE7115 | (Mazza *et al.*, 2007) |
| *Paraxis(tcf15)* | Transcription factor 15 | NV2g010865000.1 | NVE19720 | (Steinmetz *et al.*, 2017) |
| *Patch* | Protein patched homolog 1 | NV2g000986000.1 | NVE25227 | (Matus et al., 2008) |
| *Pla2* | Similar to Neutral phospholipase A2 homolog taipoxin beta chain 2 | NV2g015603000.1 | NVE16820, NVE16821 | |
| *PRGamide* | Eukaryotic translation initiation factor 3 subunit A (mis-annotated) | NV2g016299000.1 | NVE226 | (Hayakawa et al., 2019) |
| *Protocadherin* | Protocadherin-11 X-linked | NV2g002532000.1 | NVE5361, NVE11705 | |
| *Pse2b* | Similar to DELTA-alicitoxin-Pse2b | NV2g012610000.1 | NVE4235 | |
| *Pxn* | Peroxidasin | NV2g009225000.1 | NVE21612 | |

| | | | | |
|---|---|---|---|---|
| *Rgm* | Repulsive guidance molecule BMP co-receptor b | NV2g024312000.1 | NVE15735 | (Leclere and Rentzsch, 2014) |
| *RWamide.1* | Unknown | NV2g010309000.1 | NVE2302, NVE24308 | (Koch and Grimmelikhuijzen, 2020) |
| *RWamide.2* | Unknown | NV2g010311000.1 | NVE24307 | (Koch and Grimmelikhuijzen, 2020) |
| *RWamide.3* | Unknown | NV2g010312000.1 | NVE11178, NVE20010, NVE24306 | (Koch and Grimmelikhuijzen, 2020) |
| *Scgn* | Secretagogin | NV2g004529000.1 | NVE10899 | |
| *Sfrp1* | Secreted frizzled-related protein 2 | NV2g014654000.1 | NVE8371 | (Leclere et al., 2016) |
| *Six4/5* | Homeobox protein SIX4/5 | NV2g010728000.1 | NVE17554 | |
| *SnailA* | Zinc finger protein SNAI2 | NV2g000472000.1 | NVE13986 | (Fritzenwanker *et al.*, 2004) |
| *SoxB2* | Transcription factor sox-2 | NV2g004477000.1 | NVE24655 | (Richards and Rentzsch, 2014) |
| *Svep1* | Sushi, von Willebrand factor type A, EGF and pentraxin domain-containing protein 1 | NV2g003560000.1 | NVE9234, NVE19648, NVE9235, NVE9623 | |
| *Tbx15* | T-box transcription factor TBX15 | NV2g008376000.1 | NVE7505 | |
| *Tbx4/5* | T-box transcription factor TBX4/5 | NV2g016222000.1 | NVE21767 | |
| *Tgfr3* | TGF-b receptor type III | NV2g025402000.1 | NVE9533 | |
| *Thsd1* | Bone morphogenetic protein 1 | NV2g003253000.1 | NVE8009 | |
| *Thsd4* | Thrombospondin type-1 domain-containing protein 4 | NV2g022999000.1 | NVE18837, NVE5829, NVE18838 | |
| *Tlx* | T-cell leukemia homeobox protein 2 | NV2g011935000.1 | NVE21231 | |
| *Tmprss9* | Transmembrane protease serine 9 | NV2g025524000.1 | NVE16548 | |
| *tropomyosin* | Tropomyosin alpha-4 chain | NV2g008895000.1 | NVE1725 | |
| *Tspear* | Thrombospondin-type laminin G domain and EAR repeat-containing protein | NV2g008096000.1 | NVE16993 | |
| *Twist* | Twist-related protein | NV2g010864000.1 | NVE19719 | (Martindale et al., 2004) |
| *Uncx* | Homeobox protein unc-4 homolog | NV2g006849000.1 | NVE7609 | |
| *Wnt4a* | Protein Wnt-4a | NV2g022664000.1 | NVE3111 | |
| *Wnt5* | Protein Wnt-5a | NV2g003611000.1 | NVE1780, NVE20298 | |
| *Wntless* | Protein wntless homolog B | NV2g001854000.1 | NVE20439, NVE23246 | |
| *Zic3* | Zinc finger protein ZIC 3 | NV2g017755000.1 | NVE6690 | |

**References:**

Babonis, L.S., and Martindale, M.Q. (2017). PaxA, but not PaxC, is required for cnidocyte development in the sea anemone Nematostella vectensis. Evodevo *8*, 14. 10.1186/s13227-017-0077-7.

Cole, A.G., Kaul, S., Jahnel, S.M., Steger, J., Zimmerman, B., Reischl, R., Richards, G.S., Rentzsch, F., Steinmetz, P., and Technau, U. (2020). Muscle cell type diversification facilitated by extensive gene duplications. bioRxiv, 2020.2007.2019.210658. 10.1101/2020.07.19.210658.

Fritzenwanker, J.H., Saina, M., and Technau, U. (2004). Analysis of forkhead and snail expression reveals epithelial-mesenchymal transitions during embryonic and larval development of Nematostella vectensis. Dev Biol *275*, 389-402. 10.1016/j.ydbio.2004.08.014.

Genikhovich, G., and Technau, U. (2009a). Anti-acetylated tubulin antibody staining and phalloidin staining in the starlet sea anemone Nematostella vectensis. Cold Spring Harb Protoc *2009*, pdb prot5283. 10.1101/pdb.prot5283.

Genikhovich, G., and Technau, U. (2009b). In situ hybridization of starlet sea anemone (Nematostella vectensis) embryos, larvae, and polyps. Cold Spring Harb Protoc *2009*, pdb prot5282. 10.1101/pdb.prot5282.

Genikhovich, G., and Technau, U. (2009c). Induction of spawning in the starlet sea anemone Nematostella vectensis, in vitro fertilization of gametes, and dejellying of zygotes. Cold Spring Harb Protoc *2009*, pdb prot5281. 10.1101/pdb.prot5281.

Hayakawa, E., Watanabe, H., Menschaert, G., Holstein, T.W., Baggerman, G., and Schoofs, L. (2019). A combined strategy of neuropeptide prediction and tandem mass spectrometry identifies evolutionarily conserved ancient neuropeptides in the sea anemone Nematostella vectensis. PLoS One *14*, e0215185. 10.1371/journal.pone.0215185.

He, S., Del Viso, F., Chen, C.Y., Ikmi, A., Kroesen, A.E., and Gibson, M.C. (2018). An axial Hox code controls tissue segmentation and body patterning in Nematostella vectensis. Science *361*, 1377-1380. 10.1126/science.aar8384.

Ikmi, A., McKinney, S.A., Delventhal, K.M., and Gibson, M.C. (2014). TALEN and CRISPR/Cas9-mediated genome editing in the early-branching metazoan Nematostella vectensis. Nat Commun *5*, 5486. 10.1038/ncomms6486.

Koch, T.L., and Grimmelikhuijzen, C.J.P. (2020). A comparative genomics study of neuropeptide genes in the cnidarian subclasses Hexacorallia and Ceriantharia. BMC Genomics *21*, 666. 10.1186/s12864-020-06945-9.

Leclere, L., Bause, M., Sinigaglia, C., Steger, J., and Rentzsch, F. (2016). Development of the aboral domain in Nematostella requires beta-catenin and the opposing activities of Six3/6 and Frizzled5/8. Development *143*, 1766-1777. 10.1242/dev.120931.

Leclere, L., and Rentzsch, F. (2014). RGM regulates BMP-mediated secondary axis formation in the sea anemone Nematostella vectensis. Cell Rep *9*, 1921-1930. 10.1016/j.celrep.2014.11.009.

Martindale, M.Q., Pang, K., and Finnerty, J.R. (2004). Investigating the origins of triploblasty: 'mesodermal' gene expression in a diploblastic animal, the sea anemone Nematostella vectensis (phylum, Cnidaria; class, Anthozoa). Development *131*, 2463-2474. 10.1242/dev.01119.

Matus, D.Q., Magie, C.R., Pang, K., Martindale, M.Q., and Thomsen, G.H. (2008). The Hedgehog gene family of the cnidarian, Nematostella vectensis, and implications for understanding metazoan Hedgehog pathway evolution. Dev Biol *313*, 501-518. 10.1016/j.ydbio.2007.09.032.

Matus, D.Q., Pang, K., Marlow, H., Dunn, C.W., Thomsen, G.H., and Martindale, M.Q. (2006). Molecular evidence for deep evolutionary roots of bilaterality in animal development. Proc Natl Acad Sci U S A *103*, 11195-11200. 10.1073/pnas.0601257103.

Mazza, M.E., Pang, K., Martindale, M.Q., and Finnerty, J.R. (2007). Genomic organization, gene structure, and developmental expression of three clustered otx genes in the sea anemone Nematostella vectensis. J Exp Zool B Mol Dev Evol *308*, 494-506. 10.1002/jez.b.21158.

McWilliam, H., Li, W., Uludag, M., Squizzato, S., Park, Y.M., Buso, N., Cowley, A.P., and Lopez, R. (2013). Analysis Tool Web Services from the EMBL-EBI. Nucleic Acids Res *41*, W597-600. 10.1093/nar/gkt376.

Minh, B.Q., Schmidt, H.A., Chernomor, O., Schrempf, D., Woodhams, M.D., Von Haeseler, A., and Lanfear, R. (2020). IQ-TREE 2: new models and efficient methods for phylogenetic inference in the genomic era. Molecular biology and evolution *37*, 1530-1534.

Moreno-Mateos, M.A., Vejnar, C.E., Beaudoin, J.D., Fernandez, J.P., Mis, E.K., Khokha, M.K., and Giraldez, A.J. (2015). CRISPRscan: designing highly efficient sgRNAs for CRISPR-Cas9 targeting in vivo. Nat Methods *12*, 982-988. 10.1038/nmeth.3543.

Nakanishi, N., Renfer, E., Technau, U., and Rentzsch, F. (2012). Nervous systems of the sea anemone Nematostella vectensis are generated by ectoderm and endoderm and shaped by distinct mechanisms. Development *139*, 347-357. 10.1242/dev.071902.

Nguyen, L.-T., Schmidt, H.A., Von Haeseler, A., and Minh, B.Q. (2015). IQ-TREE: a fast and effective stochastic algorithm for estimating maximum-likelihood phylogenies. Molecular biology and evolution *32*, 268-274.

Nitzan, M., Karaiskos, N., Friedman, N., and Rajewsky, N. (2019). Gene expression cartography. Nature *576*, 132-137. 10.1038/s41586-019-1773-3.

Rentzsch, F., Anton, R., Saina, M., Hammerschmidt, M., Holstein, T.W., and Technau, U. (2006). Asymmetric expression of the BMP antagonists chordin and gremlin in the sea anemone Nematostella vectensis: implications for the evolution of axial patterning. Dev Biol *296*, 375-387. 10.1016/j.ydbio.2006.06.003.

Richards, G.S., and Rentzsch, F. (2014). Transgenic analysis of a SoxB gene reveals neural progenitor cells in the cnidarian Nematostella vectensis. Development *141*, 4681-4689. 10.1242/dev.112029.

Ryan, J.F., Mazza, M.E., Pang, K., Matus, D.Q., Baxevanis, A.D., Martindale, M.Q., and Finnerty, J.R. (2007). Pre-bilaterian origins of the Hox cluster and the Hox code: evidence from the sea anemone, Nematostella vectensis. PLoS One *2*, e153. 10.1371/journal.pone.0000153.

Saina, M., Genikhovich, G., Renfer, E., and Technau, U. (2009). BMPs and chordin regulate patterning of the directive axis in a sea anemone. Proc Natl Acad Sci U S A *106*, 18592-18597. 10.1073/pnas.0900151106.

Scholz, C.B., and Technau, U. (2003). The ancestral role of Brachyury: expression of NemBra1 in the basal cnidarian Nematostella vectensis (Anthozoa). Dev Genes Evol *212*, 563-570. 10.1007/s00427-002-0272-x.

Shinzato, C., Khalturin, K., Inoue, J., Zayasu, Y., Kanda, M., Kawamitsu, M., Yoshioka, Y., Yamashita, H., Suzuki, G., and Satoh, N. (2021). Eighteen Coral Genomes Reveal the Evolutionary Origin of Acropora Strategies to Accommodate Environmental Changes. Mol Biol Evol *38*, 16-30. 10.1093/molbev/msaa216.

Shumaker, A., Putnam, H.M., Qiu, H., Price, D.C., Zelzion, E., Harel, A., Wagner, N.E., Gates, R.D., Yoon, H.S., and Bhattacharya, D. (2019). Genome analysis of the rice coral Montipora capitata. Sci Rep *9*, 2571. 10.1038/s41598-019-39274-3.

Sievers, F., Wilm, A., Dineen, D., Gibson, T.J., Karplus, K., Li, W., Lopez, R., McWilliam, H., Remmert, M., Soding, J., et al. (2011). Fast, scalable generation of high-quality protein multiple sequence alignments using Clustal Omega. Mol Syst Biol *7*, 539. 10.1038/msb.2011.75.

Stefanik, D.J., Friedman, L.E., and Finnerty, J.R. (2013). Collecting, rearing, spawning and inducing regeneration of the starlet sea anemone, Nematostella vectensis. Nat Protoc *8*, 916-923. 10.1038/nprot.2013.044.

Steinmetz, P.R.H., Aman, A., Kraus, J.E.M., and Technau, U. (2017). Gut-like ectodermal tissue in a sea anemone challenges germ layer homology. Nat Ecol Evol *1*, 1535-1542. 10.1038/s41559-017-0285-5.

Watanabe, H., Kuhn, A., Fushiki, M., Agata, K., Ozbek, S., Fujisawa, T., and Holstein, T.W. (2014). Sequential actions of beta-catenin and Bmp pattern the oral nerve net in Nematostella vectensis. Nat Commun *5*, 5536. 10.1038/ncomms6536.

Wijesena, N., Simmons, D.K., and Martindale, M.Q. (2017). Antagonistic BMP-cWNT signaling in the cnidarian Nematostella vectensis reveals insight into the evolution of mesoderm. Proc Natl Acad Sci U S A *114*, E5608-E5615. 10.1073/pnas.1701607114.

Wolenski, F.S., Layden, M.J., Martindale, M.Q., Gilmore, T.D., and Finnerty, J.R. (2013). Characterizing the spatiotemporal expression of RNAs and proteins in the starlet sea anemone, Nematostella vectensis. Nat Protoc *8*, 900-915. 10.1038/nprot.2013.014.
